# Xist deletion in T cells protects against influenza-induced lung injury

**DOI:** 10.64898/2026.11.20.768059

**Authors:** Nuriban Valero-Pacheco, Katherine S. Forsyth, Natalie Toothacre, Robert T. Trudeau, Emma Welter, Nikhil Jiwrajka, Sonresa Molina-Vidales, Camille M. Syrett, Bomyi Lim, Montserrat C. Anguera

## Abstract

Sex differences in antiviral immunity are well established, yet the contribution of X-Chromosome Inactivation (XCI) to T cell responses during infection is unknown. Here, we identify an unexpected role for Xist RNA in shaping T cell immunity and pulmonary injury following influenza infection. We show that the inactive X chromosome (Xi) in pulmonary CD4+ and CD8+ T cells undergoes dynamic epigenetic remodeling, with limited Xist RNA localization, differential enrichment of H3K27me3 and H2AK119Ub modifications, and activation-dependent XCI escape of immune-related genes, including *Il2rg and Cxcr3*. Remarkably, T cell-specific *Xist* deletion protects female mice from influenza-induced weight loss and pulmonary pathology despite comparable early viral burdens. Transcriptomic profiling reveals increased expression of X-linked escape genes and antiviral and interferon-responsive genes, particularly in CD8+ T cells. Furthermore, Xist-deficient CD4^+^ and CD8^+^ antigen-experienced (CD44^+^CD11a^+^) T cells exhibit increased biallelic *Cxcr3* expression, elevated pulmonary Cxcr3 protein levels, and enhanced IFN-γ and TNF responses following *ex vivo* stimulation. Together, our findings reveal a previously unrecognized role for Xist RNA in regulating female antiviral T cell responses and demonstrate that enhanced T cell effector potential can accompany reduced infection-associated pulmonary injury.

## INTRODUCTION

Females mount stronger adaptive responses to most viral infections than males, including influenza A virus^1^. These sex differences manifests in various ways, including faster viral clearance, reduced severe immunopathology, and lower rates of post-infection morbidity^2^. The biological origins for sex differences with immune responses are the sex hormones and sex chromosomes, both of which likely contribute to sex-dependent influenza responses. Notably, incidence data associated with influenza infections shows a strong sex bias among pre-pubescent children, with females being at lower risk of severe disease outcomes than males^3^, suggesting that genetic origins from the sex chromosomes may also contribute to sex differences with viral pathogenesis and immunity after influenza infection. The mammalian X chromosome contains many conserved immune-related X-linked genes including *FOXP3, CD40LG, IL2RG*, and *CXCR3*^4^, suggesting that X-linked gene dosage and expression can be an important contributor to female-biased immune responses to influenza infections.

In female mammals, one X chromosome is randomly selected for transcriptional silencing in a process called X-Chromosome Inactivation (XCI) that equalizes X-linked gene expression between XX and XY individuals^5^. XCI is initiated and maintained by the long noncoding RNA Xist, which is expressed and localized to the inactive X (Xi)^6^. Xist RNA can recruit Polycomb Repressive Complexes (PRC1, PRC2) that deposit repressive histone tail modifications H2AK119-ubiquitin (Ub) and H3K27me3 across the Xi, and both Xist RNA and these heterochromatic modifications can be visualized cytologically in most female somatic cells^7,8^. Xist RNA and heterochromatic modifications function to maintain transcriptional silencing across most of the Xi. However, there are X-linked genes that ‘escape’ XCI and are expressed from the Xi, including *Xist*, and biallelic expression of these X-linked genes can result in sex-biased expression^9^. Constitutive XCI escape genes have a Y-linked homolog and are expressed in diverse cell types^10^, and some X-linked genes escape XCI in specific cell contexts, such as *TLR7* in immune cells^11,12^.

Unlike most female somatic cells, immune cells maintain XCI differently and the Xi is less heterochromatic and more relaxed yet remains dosage compensated^13–15^. Unstimulated T cells lack cytological enrichment of Xist RNA ‘clouds’ and foci of H3K27me3 and H2AK119Ub at the Xi^16,17^. *In vitro* stimulation through activation of the T cell receptor (TCR), and subsequent NF-kB activation and nuclear localization, is required to localize Xist RNA and heterochromatic marks to the Xi^15^. The atypical XCI maintenance features in female lymphocytes may allow conditional XCI escape of immune-relevant X-linked genes, without full Xi reactivation^15,18^. Unstimulated T cells exhibit expression of X-linked immunity genes, such as *Il2rg* from the Xi regardless of TCR activation *in vitro*, while other X-linked immunity genes, such as *Cxcr3*, exhibit XCI escape in *in vitro* activated T cells^15^. It is unknown whether the epigenetic features of XCI maintenance are dynamically remodeled in T cells in response to a viral infection *in vivo*, and whether Xist RNA is required to preserve X-linked gene dosage for optimal antiviral T cell function.

Lung-infiltrating CD4^+^ and CD8^+^ T cells are essential for influenza clearance and differentiation into effector T cell subsets capable of cytokine production. Recruitment of CD4^+^ and CD8^+^ T cells to the infected lung depends on TCR activation and chemokine signaling, including from X-linked Cxcr3-mediated trafficking^19^. *CXCR3*/*Cxcr3* can be biallelically expressed in activated human and mouse T cells, and *Leishmania* infection increases numbers of biallelic *Cxcr3* T cells ^15,17,20^. Intriguingly, biallelic *Cxcr3*-expressing T cells from infected mice exhibit increased effector function^20^, which suggests that other XCI escape genes may influence T cell antiviral responses in a sex-specific manner. It is unknown how viral infections affect gene expression from Xi in T cells, and whether XCI escape can influence T cell function and immune responses to viral infection.

Here, we investigated how influenza infection influences XCI maintenance and whether Xist RNA regulates antiviral T cell responses *in vivo*. We found that pulmonary CD4^+^ and CD8^+^ T cells exhibit dynamic changes in Xist RNA localization and epigenetic features of the Xi during infection. Allele-specific epigenomic and transcriptional profiling revealed preferential enrichment of H3K27me3, but not H2AK119Ub, on the Xi, together with activation-dependent escape from XCI of immune-related genes, including *Cybb*, *Il2rg*, and *Cxcr3*. Unexpectedly, T cell-specific deletion of *Xist* protected female mice from influenza-induced weight loss and pulmonary injury, despite comparable early viral burdens. Xist-deficient T cells exhibited increased expression of antiviral and interferon-stimulated genes, particularly in CD8^+^ T cells, accompanied by increased biallelic *Cxcr3* expression and elevated Cxcr3 protein levels. Moreover, CD4^+^ and CD8^+^ CD44^+^CD11a^+^ antigen-experienced (AgExp) T cells from infected *Xist* deleted mice displayed enhanced IFN-γ and TNF responses following *ex vivo* stimulation. Together, these findings identify Xist RNA as a previously unrecognized regulator of female antiviral T cell responses and reveal that increased T cell effector potential can accompany reduced infection-associated pulmonary pathology, establishing a link between XCI maintenance and antiviral immunity.

## RESULTS

### Influenza infection results in variable Xist RNA and H2AK119Ub enrichment on the Xi in lung CD4^+^ and CD8^+^ naive and antigen-experienced T-cell subsets

We infected wildtype (WT) female mice with influenza A virus PR8 and investigated the timing for the appearance of Xist RNA localization and the heterochromatic histone tail modification H2AK119Ub, both of which are usually cytologically enriched on the Xi in female somatic cells, missing in unstimulated T cells, and appear at the Xi following T cell receptor engagement *in vitro*^15^. We sorted lung naïve (CD44^-^CD11a^-^; ‘Naive’) and antigen-experienced (CD44^+^CD11a^+^; ‘AgExp’) CD4+ or CD8+ T cell subsets at either 5 or 10 days post-infection (dpi), which correspond to a timepoint of early recruitment of effector T cells to the infected lung and high lung viral burdens (5 dpi) or a late effector time point characterized by near-peak pulmonary T cell responses coincident with viral clearance (10 dpi)^21–23^. We performed sequential Xist RNA FISH and immunofluorescence (IF) for H2AK119Ub on these T cell subsets isolated from the lung, mediastinal draining lymph nodes (dLN), and spleen from infected mice, which revealed a mixture of cells with bright canonical Xist RNA ‘clouds’ and dispersed pinpoints, yet the majority of cells lacked Xist RNA signals (**Figure 1, Supplemental Figure 1**). In the lung, we observed more nuclei with Xist RNA signals in both CD4^+^ and CD8^+^ T cells at 5 dpi compared to 10 dpi (**Fig. 1A-D**), and CD4^+^ AgExp T cells exhibited more Xist RNA signals compared to CD4^+^ naive T cells (**Figure 1E**). Yet in the dLN, Xist RNA signals were comparable across 5 and 10 dpi for both CD4^+^ and CD8^+^ T-cell subsets (**Supplemental Figure 1A-D**), where 30-40% of nuclei contained Xist RNA clouds or pinpoints (**Figure S1E**). Examination of Xist RNA patterns in spleens of infected female mice revealed fewer nuclei with Xist RNA signals (**Figure S2A-D**), with no significant differences between Naive and AgExp T cells (**Figure S2E**). Thus, for both CD4+ and CD8+ T cells, location and time post-viral challenge appears to correlate more directly with Xist RNA signals than activation status, suggesting that antigen availability to stimulate the T cell receptor is critical *in vivo*, similar to splenic CD3+ T cells stimulated with anti-CD3/28 *in vitro*^15^.

**Figure 1.**
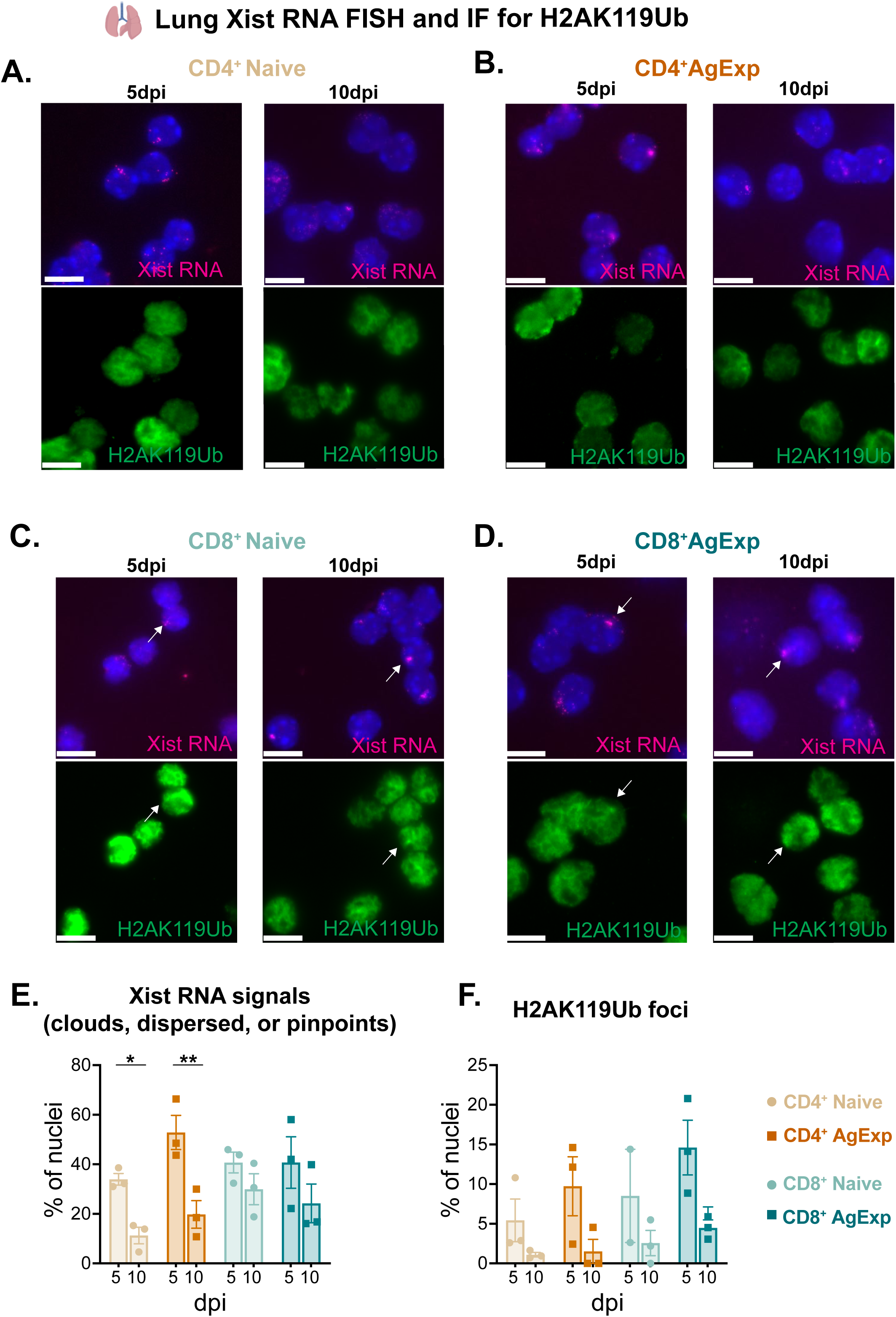
Flu-infected lung Naive and AgExp T cells exhibit more Xist RNA clouds at 5 vs 10 dpi with few H2AK119Ub foci. Lung CD4^+^ and CD8^+^ naive (CD44^-^CD11a^-^) and antigen-experienced (AgExp, CD44^+^CD11a^+^) T cells were sorted from influenza PR8-infected female C57BL/6 mice at 5 and 10 days post-infection (dpi) and analyzed by sequential Xist RNA FISH and H2AK119Ub immunofluorescence (IF). Representative images of CD4^+^ Naive (**A**), CD4^+^ AgExp (**B**), CD8^+^ Naive (**C**), and CD8^+^ AgExp (**D**) T cells are shown. Arrows indicate representative Xist RNA signals and corresponding H2AK119Ub foci. Scale bars, 10 μm. Quantification of the percentage of nuclei with Xist RNA signals (**E**) and H2AK119Ub foci (**F**) is shown for each T-cell subset. For each biological replicate and T-cell subset, 99–215 nuclei were evaluated for Xist RNA and 41–164 nuclei for H2AK119Ub IF. At 5 dpi, each data point represents a pool of n = 2 mice; at 10 dpi, each data point represents one mouse. Data are shown as mean ± SEM. CD4^+^ and CD8^+^ T-cell subsets were analyzed separately by ordinary two-way ANOVA with time point and T-cell subset as factors, followed by Holm–Šídák multiple-comparisons tests comparing 5 vs. 10 dpi within each T-cell subset. \**P* < 0.05,\*\**P* < 0.01.

IF detection for nuclear H2AK119Ub localization, which is cytologically enriched at the Xi in most female somatic cells and *in vitro* activated bulk CD3^+^ T cells, revealed that there were few CD4^+^ or CD8^+^ T cells with an H2AK119Ub focus following viral infection (**Fig.1, Fig S1, S2).** In the lung, the 5dpi CD4^+^ and CD8^+^ T cells trended toward higher frequencies of H2AK119Ub foci ranging between 5-15% of nuclei, but the differences were not statistically significant (**Fig. 1F**). We did observe infrequent co-localization of bright Xist RNA signals with H2AK119Ub foci in T cells across tissues (white arrows, **Fig. 1, Fig. S1-2).** There were fewer H2AK119Ub foci in T cells from dLNs and spleens (5-10%) compared to lung T cells, irrespective of infection timepoint (**Fig. S1F, S2F**). Thus, flu infection does not consistently result in cytological enrichment of Xist RNA and H2K119Ub at the Xi in CD4^+^ and CD8^+^ T-cell subsets across lung and lymphoid tissues at 5-10 dpi.

### The Xi has low levels of H2AK119Ub yet is enriched with H3K27me3 after influenza infection in Naive and AgExp lung T-cell subsets

Because flu infection resulted in few T cells containing H2AK119Ub foci, we investigated the enrichment of heterochromatic modifications H3K27me3 and H2AK119Ub at higher resolution across the Xi in lung CD4^+^ and CD8^+^ T cells after infection. We utilized a F1 hybrid mouse model of skewed XCI where female C57BL/6 mice (‘mus’) containing a heterozygous *Xist* deletion were mated to male *Mus castaneus (‘*cast*’)* mice (‘F1 mus x cast’), and single nucleotide polymorphisms can distinguish the *mus* and *cast* genomes (**Figure 2A**)^15,18^. We infected female F1 mus x cast mice with influenza, then sorted CD4^+^ and CD8^+^ Naive and AgExp T cells from lungs at 5 and 10 dpi and performed allele-specific CUT&RUN for H2AK119Ub and H3K27me3 histone marks (**Fig. 2A**). Surprisingly, quantification of total CPM normalized reads for H2AK119Ub indicate similar levels for Xi and Xa, with low levels of H2AK119Ub across all cells and timepoints (**Fig. 2B**). We found that the Xi in both CD4^+^ and CD8^+^ Naive T cells had even less H2AK119Ub enrichment compared to the Xa (**Fig. 2B**), yet there were similar levels of H2AK119Ub for Xi and Xa in AgExp T cells (**Fig. 2B**). For H3K27me3, the Xi contained significantly higher levels compared to the Xa across all cell types and timepoints with slightly lower levels at 10 compared to 5 dpi of CD4+ or CD8+ T cells (**Fig. 2C**). H2AK119Ub is evenly distributed across the Xi in CD4^+^ Naive and AgExp T cells between 5-10 dpi (**Fig. 2D,E**), yet d-score analyses indicate increased H2AK119Ub accumulation at gene and promoter (‘genic’) regions of CD4^+^ Naive T cells for 10 dpi (**Fig. 2F,G**). H3K27me3 marks become reduced across various regions of the Xi from 5-10 dpi (**Fig. 2H**), and d-score analyses show significant H3K27me3 reductions at both genic and intergenic regions (**Fig. 2I**). While CD8^+^ Naive T cells exhibited small changes with H2AK119Ub and H3K27me3 across Xi between 5-10 dpi (**Fig. 2J,K**), the CD8^+^ AgExp T cells have regions that lost both marks (**Fig. 2L,M**). D-score analyses revealed that H2AK119Ub marks significantly accumulate at genic and intergenic regions in 10 dpi CD8^+^ Naive T cells (**Fig. 2N**), yet H3K27me3 decreases at intergenic regions (**Fig. 2O**). For CD8^+^ AgExp T cells, H2AK119Ub marks accumulate in 10 dpi at both genic and intergenic regions (**Fig. 2P**), while H3K27me3 marks significantly decrease at genic and intergenic regions of 10 dpi (**Fig. 2Q**). Examination of the *Hox* locus revealed expected enrichment patterns of both H3K27me3 and H2AK119Ub at silenced genes (**Fig. S3A**) and allelic distribution of both modifications for chromosome 13 were comparable (**Fig. S3B**) for both T cell subsets at 5-10 dpi. D-score analyses for genic and intergenic enrichment of H2AK119Ub and H3K27me3 were mostly similar across chromosome 13 for Naive and AgExp T-cell subsets (**Fig. S3C-I**). Enrichment of H2AK119Ub and H3K27me3 marks were mostly correlated across the Xi for 5 and 10 dpi in T-cell subsets (**Fig. S3J-M),** except at distal regions (∼160 Mb) in Naive T cells where there was less H2AK119Ub compared to AgExp T cells. Thus, at the Xi across infected lung T cell subsets is enriched for H3K27me3, but not H2AK119Ub, at both timepoints with high lung viral burden (5 dpi) and at the near-peak pulmonary T cell response coincident with viral clearance (10 dpi).

**Figure 2.**
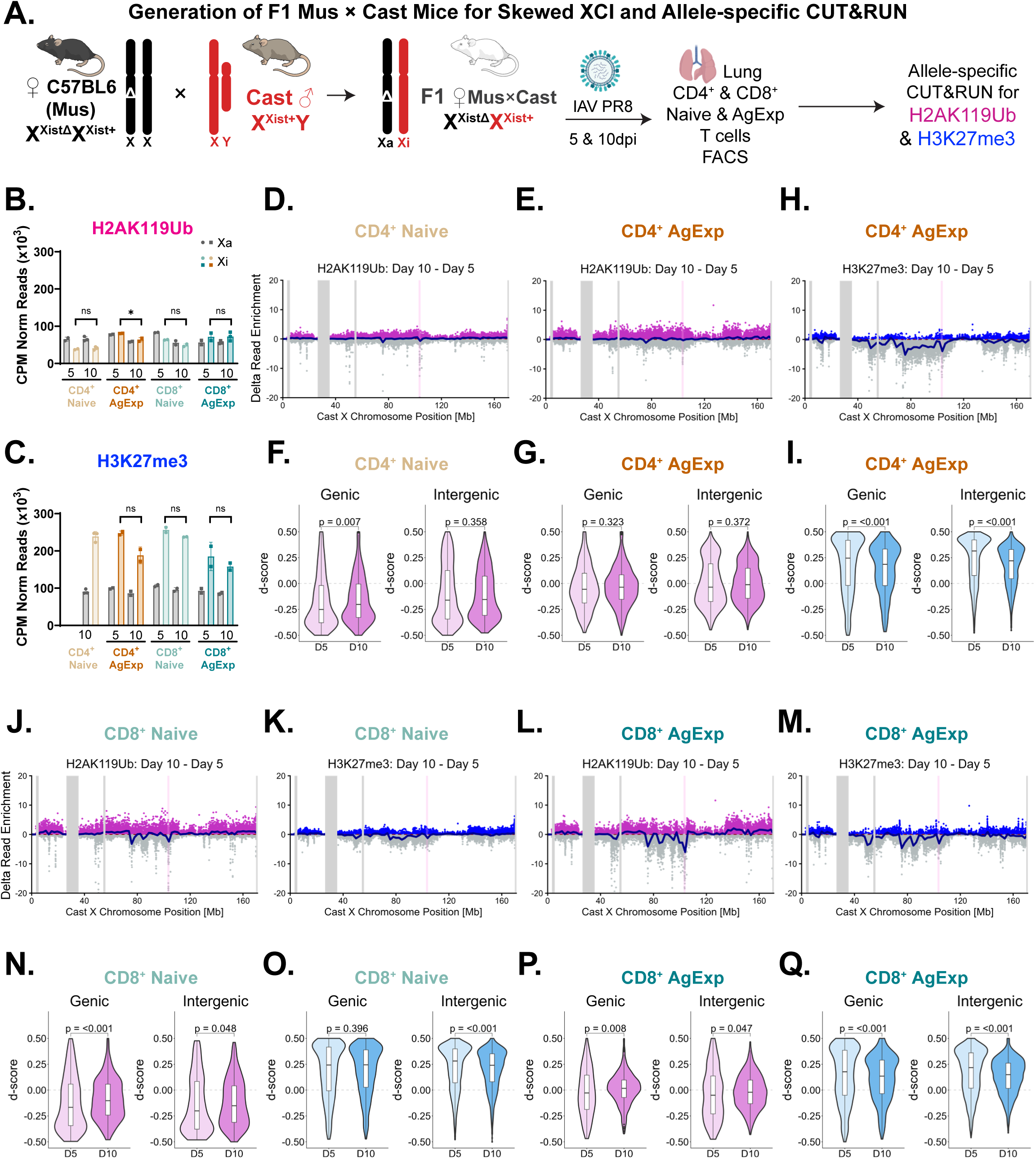
Allele-specific epigenomic profiling of H3K27me3 and H2AK119Ub across the X chromosomes in the pulmonary T cell subsets from flu-infected mice. (**A**) Experimental design for allele-specific CUT&RUN analysis of heterochromatic marks from female F1 mus x cast mice. The paternal Cast-derived X chromosome is Xi and the maternal BL6-derived X chromosome is Xa. Lung CD4^+^ and CD8^+^ Naive and AgExp T cells were isolated by FACS at 5 and 10 dpi. (**B-C**) Sum of CPM normalized reads across the Xa (black) and Xi (colors) in lung CD4^+^ Naive, CD4^+^ AgExp, CD8^+^ Naive, and CD8^+^ AgExp T cells at 5 and 10 dpi for H2AK119Ub (**B**) and H3K27me3 (**C**). Unpaired t-test performed on Xi reads between 5 and 10 dpi within each cell type, * *P* < 0.05. (**D-E**) Delta accumulation of H2AK119Ub between 5 and 10 dpi at 10 kb bins (dots) across the Xi in CD4^+^ Naive (**D**) and AgExp (**E**) T cells. Positive accumulation is in purple and negative is in gray. The blue line represents the locally estimated scatterplot smoothing (LOESS) regression across all 10 kb bins. Vertical gray bars are unmappable regions. Vertical pink bar is the *Xist* locus. (**F-G**) Mean d-score [(Xireads/(Xireads+Xareads)) – 0.5] at genic and intergenic regions between 5 and 10 dpi in CD4^+^ Naive T cells. Genic regions are defined as +1 kb from transcription start site (TSS) to the transcription end site (TES) and intergenic regions are defined as 10 kb bins not overlapping genes or their promoters. Adjusted *P* values calculated using a Kruskal-Wallis test followed by a Dunn’s test with Benjamini-Hochberg correction for pairwise comparisons. (**H**) Delta accumulation of H3K27me3 between 5 and 10 dpi at 10 kb bins (dots) across the Xi in CD4^+^ AgExp T cells. (**I**) Mean d-score analyses for H3K27me3 enrichment in CD4^+^ AgExp T cells. (**H-I**) As D-E for H3K27me3 in CD4^+^ AgExp T cells. (**J-K**) Delta accumulation of H2AK119Ub (J) and H3K27me3 (K) between 5 and 10 dpi in CD8^+^ Naive T cells. (**L-M**) Delta accumulation of H2AK119Ub (L) and H3K27me3 (M) between 5 and 10 dpi in CD8^+^ AgExp T cells. (**N-O**) Mean d-score analyses for H2AK119Ub (N) H3K27me3 (O) enrichment in CD8^+^ Naive T cells. (**P-Q**) Mean d-score analyses for H2AK119Ub (P) H3K27me3 (Q) enrichment in CD8^+^ AgExp T cells.

Next, we asked how activation status of T cells during the near-peak T cell pulmonary response (10 dpi) effected Xi enrichment of marks by comparing lung-derived AgExp and Naïve cells. We determined the delta read enrichment for each heterochromatic mark for AgExp cells relative to Naïve cells. For CD4+ AgExp T cells, we found that there is more enrichment of H2AK119Ub compared to Naïve T cells across the Xi (**Fig. 3A**), yet H3K27me3 levels are variably enriched between AgExp and Naïve cells at different regions across the Xi (**Fig. 3B**). Examining accumulation differences specifically at Xi genes, we observed that 149 Xi-linked genes are significantly enriched for H2AK119Ub marks and 10 Xi-linked genes are depleted for H2AK119Ub in CD4 AgExp cells (**Fig. 3C**). For H3K27me3, there were 16 Xi-linked genes that are significantly enriched and 196 Xi-linked genes that are depleted for this mark in CD4+ T cells (**Fig. 3D**). Examining heterochromatic mark changes across the Xi for CD8+ AgExp T cells, we observed the same trends as CD4+ T cells, with a smaller magnitude of changes at discreet X-linked regions (**Fig. 3E, F**). CD8+ AgExp T cells significantly gained H2AK119Ub at 67 Xi genes and lost this mark at 5 Xi genes (**Fig. 3G**). The H3K27me3 marks were significantly reduced in 225 Xi-linked genes and accumulated at 10 Xi-linked genes in CD8+ AgExp T cells (**Fig. 3H**). Thus, the Xi in infected lung Naïve and AgExp CD4+ and CD8+ T cells is mostly enriched with H3K27me3 but not H2AK119Ub (**Fig.2C**), and AgExp T cells exhibit activation state dependent changes with enrichment of both H2AK119Ub and H3K27me3 marks compared to Naïve T cells.

**Figure 3.**
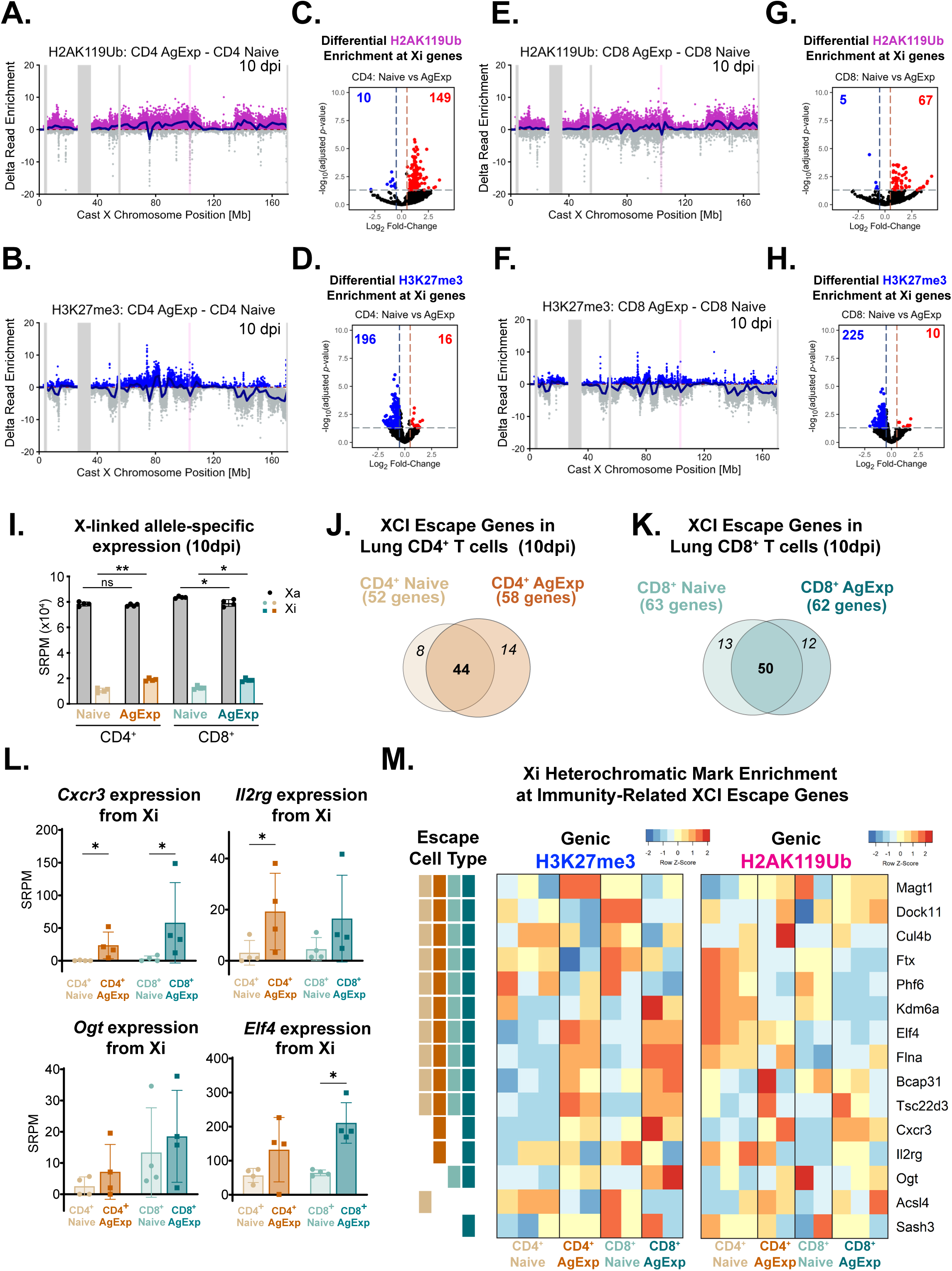
CD4^+^ and CD8^+^ AgExp T cells show enrichment changes for both H2AK119Ub and H3K27me3 marks and exhibit infection-specific XCI escape genes. (**A-B**) Delta accumulation of H2AK119Ub (A) and H3K27me3 (B) between lung CD4^+^ Naive and CD4^+^ AgExp T cells at 10 kb bins (dots) across the Xi at 10 dpi. Positive accumulation is in purple (A) or blue (B) and negative is in gray. The blue line represents the LOESS regression across all 10 kb bins. Vertical gray bars are unmappable regions. Vertical pink bar is the *Xist* locus. (**C-D**) Differential enrichment analysis of H2AK119Ub (C) and H3K27me3 (D) at X-linked genes between lung CD4^+^ Naive and CD4^+^ AgExp T cells at 10 dpi. Vertical lines represent a ±1.5 fold-change in mark enrichment and horizontal line represents an adjusted *P* value of 0.05. Genes with significant enrichment in CD4^+^ AgExp as compared to CD4^+^ Naive T cells are colored red, and those depleted are colored blue. (**E-H**) Same analyses as A-D for lung CD8^+^ Naive and CD8^+^ AgExp T cells at 10 dpi. (**I**) Allele-specific expression [SNP-reads per million (SRPM)] from the Xi (black) and Xa (colors) in CD4^+^ and CD8^+^ Naive and AgExp T cells isolated from the lung 10 dpi analyzed by ordinary two-way ANOVA followed by Holm–Šídák multiple-comparisons tests \**P* < 0.05,\*\**P* < 0.01. (**J-K**) Overlap of identified escape genes between CD4^+^ (J) and CD8^+^ (K) Naive and AgExp in T cells from the lung 10 dpi. (**L**) Xi-specific gene expression (SPRM) from select immunity-related escape genes. *P*-values calculated using a Kruskal-Wallis test followed by Dunn’s multiple comparisons test, \**P* < 0.05. (**M**) Normalized Xi enrichment of H2AK119Ub and H3K27me3 at immunity-related escape genes identified in CD4^+^ and CD8^+^ Naive and AgExp T cells from the lung 10 dpi [+1 kb from transcription start site (TSS) to the transcription end site (TES)]. Each column represents 1 biological replicate. The cell type(s) in which escape has been identified is denoted by colored boxes on the left.

### T-cell subsets from flu-infected lungs have a Xi that is dosage compensated yet exhibit cell type-specific XCI escape genes

Because we observed dynamic enrichment of repressive epigenetic modifications across the Xi between Naïve and AgExp T cells, we examined allele specific X-linked gene expression patterns from T-cell subsets from lungs of infected mice. We repeated influenza PR8 infections of female F1 mus x cast mice, and collected CD4^+^ and CD8^+^ Naive and AgExp T cells from 10 dpi infected lungs for allele-specific gene expression profiling. Principal component analyses (PCA) indicate that genome-wide transcriptional profiles for replicates of each T-cell subset clusters together, with Naive and AgExp T cells distinguished in PC1, and CD4^+^ vs CD8^+^ cells in PC2 axis (**Fig. S4A**). Unsupervised clustering of genome-wide differentially expressed genes (DEGs) further distinguished Naive and AgExp T cells, consistent with their sorting phenotypes, with increased expression of effector-associated genes, including *Nkg7*, *Ccr2*, and *Cd44*, in AgExp T cells, and Naive-associated genes, such as *Tcf7*, *Ccr7*, and *Sell*, in Naive T cells (**Fig. S4B; Supplemental File 1**). We found that the Xi was dosage compensated with less transcription compared to the Xa across all four T-cell subsets, and the Xi in AgExp T cells had significantly more expression compared to Naive T cells (**Fig. 3I**). Allele-specific expression for all autosomal genes were similar between maternal (C57BL/6) and paternal (cast) alleles across T-cell subsets (**Fig. S4C**). Comparison of Xi expressed genes, which ‘escape’ XCI, between CD4^+^ Naive and AgExp T cells revealed that 44 XCI genes are shared between cellular states, and there are unique XCI escapees specific to Naive (8 genes) and AgExp (14 genes) T cells (**Fig. 3J; Supplemental File 2**). Lung CD8^+^ T cells from infected female mice exhibited more XCI escape genes compared to CD4^+^ T cells, and 50 Xi genes were expressed in both CD8^+^ Naive and AgExp T cells (**Fig. 3K; Supplemental File 2**). We found that some T-cell immunity-related X-linked genes escape XCI in T cell subsets from infected lungs, notably *Elf4, Il2rg*, *Tsc22d3, Magt1, Phf6, Ogt, Cybb,* and *Cxcr3,* in addition to the constitutive XCI escape genes *Kdm6a*, *Ddx3x* and *Eif2s3x* (**Supplemental File 2**). Notably, we found that X-linked genes important for Th1 cellular responses, *Il2rg* and *Cxcr3,* both escape XCI in CD4^+^ and CD8^+^ AgExp T cells (**Fig. 3L**). We also observed expression of novel XCI escape genes *Phf6*, *Rbm3,* and *Tktl1* in Naive and AgExp T cells that have not been previously reported to escape XCI in splenic bulk CD3^+^ T cells stimulated *in vitro*^15^. *Xist* was robustly expressed across all T-cell subsets, and AgExp T cells had significantly higher expression compared to Naive T cells (**Fig. S4D**), likely reflecting increased transcriptional activity associated with lymphocyte activation. Together, we observed that, for all subsets, dosage compensation was largely intact, yet some X-linked genes with roles in immunity and T cell function escape in some or all T cell subsets examined.

Next, we asked how H3K27me3 and H2AK119Ub enrichment levels change at XCI gene regions (including promoters) of all infection-specific T cell XCI escape genes, and 15 immunity-related XCI escape genes across Naive and AgExp T cells. We found that there is variability with heterochromatic mark changes comparing Naive to AgExp T cells across the infection-specific T-cell XCI escape genes, including X-linked immunity-related genes (**Fig. S4E**). For example, the *Il2rg, Dock11, Phf6, Ftx,* and *Cul4b* regions lose some H3K27me3 and H2AK119Ub modifications in AgExp compared to Naive T cells (**Fig. 3M**). The *Ogt, Elf4*, and *Flna* regions gain just H3K27me3, but not H2AK119Ub marks in AgExp T cells (**Fig. 3M**). Finally, the *Tsc22d3, Bcap31, and Cxcr3* regions gain both H3K27me3 and H2AK119Ub modifications in AgExp compared to Naive T cells (**Fig. 3M**). Taken together, some T-cell specific genes with known antiviral immune function escape XCI in T cell subsets from flu-infected lungs and exhibit variable patterns of heterochromatic mark enrichment across genic regions in AgExp compared to Naive T cells.

### Conditional deletion of Xist in T cells protects female mice from flu-induced weight loss and lung injury

Because flu infection altered the expression of some Xi genes in lung AgExp T cells, we asked whether perturbations with XCI maintenance would impact T cell-mediated responses following infection. We used CD4-Cre Recombinase mice to delete *Xist* in CD4^+^ and CD8^+^ T cells on both C57BL/6 and BALB/c backgrounds. CD4-Cre Recombinase is expressed in thymocytes starting at the double-positive stage and continuously in single-positive CD4^+^ and CD8^+^ T cells^24^, after the maintenance phase of XCI has been established (**Figure 4A**). Xist ^2lox/2lox^ mice were mated to CD4-Cre Recombinase animals, and heterozygous Xist ^2lox/+^;CD4^+/-^ pups were crossed to generate Xist ^2lox/2lox^ CD4^+/-^ (‘Xist TcKO’) female mice. Using sequential Xist RNA FISH and IF for H3K27me3, we confirmed that *Xist* was deleted in 100% of splenic CD3^+^ T cells from Xist TcKO female mice (**Fig. S5A-B**). Female TcKO mice are fertile and viable, with comparable survival rates as wildtype (Xist WT) female mice (**Figure S5C**).

**Figure 4.**
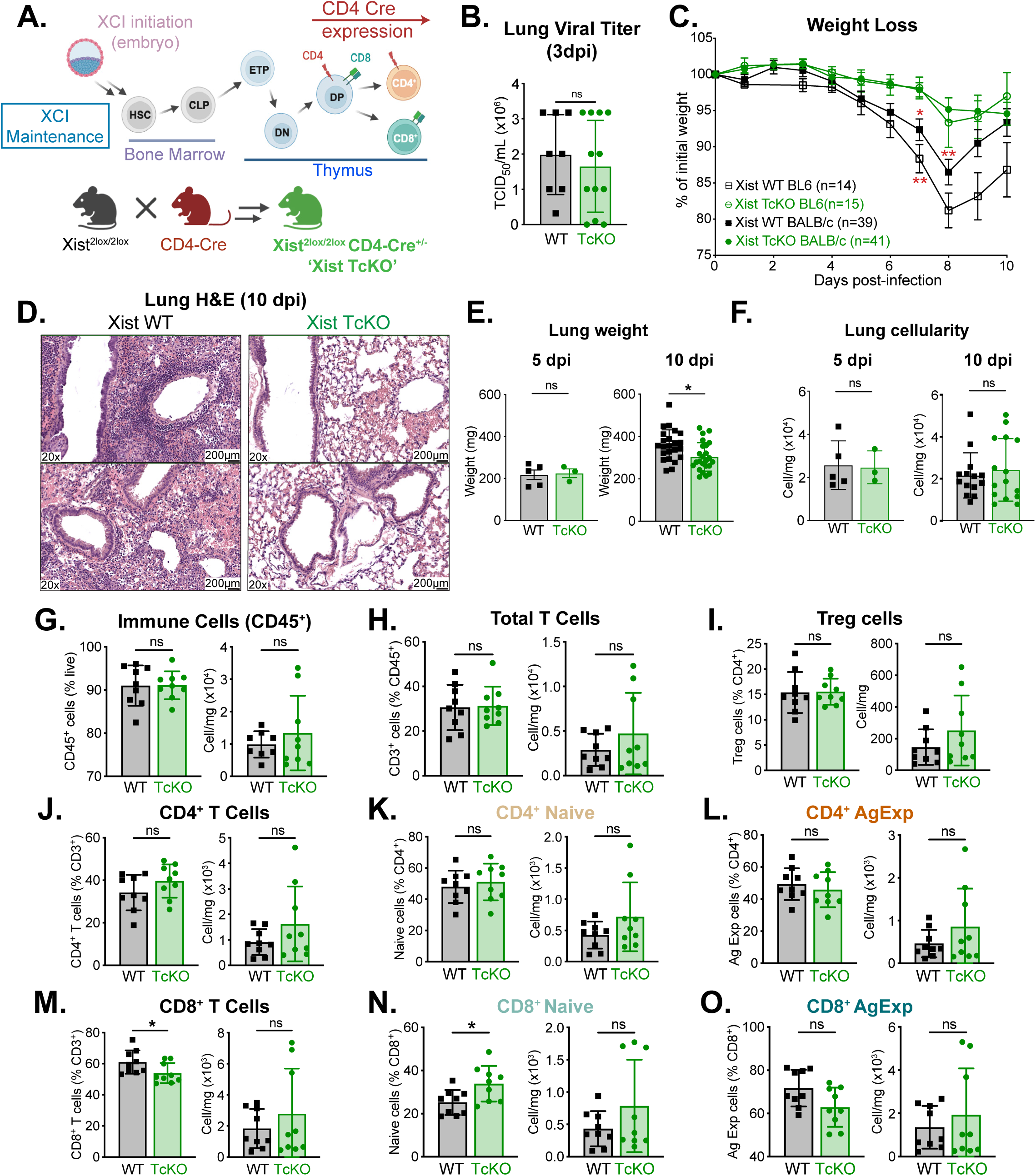
T cell-specific *Xist* deletion protects female mice from influenza-induced lung pathology. (**A**) Schematic of the Xist T cell conditional knockout (TcKO) model generated by crossing Xist2lox mice with CD4-Cre mice. CD4-Cre-mediated deletion occurs after XCI initiation and during thymocyte development. (**B**) Infectious viral titers in lung homogenates at 3 dpi, determined by TCID50 assay in MDCK cells. Data were pooled from three independent experiments with 2–5 mice per group and are shown as mean ± SD. (**C**) Body weight change following influenza PR8 infection in Xist WT and Xist TcKO mice on C57BL/6 and BALB/c backgrounds, expressed as percentage of initial body weight. BALB/c data were pooled from ten independent experiments with 3–7 mice per group per experiment, and C57BL/6 data were pooled from four independent experiments with 3–5 mice per group. Longitudinal weight-loss data for each genetic background were analyzed separately using a mixed-effects model (REML) followed by Šídák’s multiple-comparisons tests at each time point and are shown as mean ± SEM. (**D**) Representative H&E-stained lung sections from Xist WT and Xist TcKO mice at 10 dpi, representative of n = 4 WT and n = 7 TcKO mice. Scale bars, 200 μm. (**E**) Lung weights at 5 and 10 dpi. Data for 10 dpi were pooled from six independent experiments with 2–5 mice per group. Each data point represents one mouse, and Xist WT and Xist TcKO groups were compared using two-tailed Mann–Whitney tests. Data are shown as mean ± SD (**F**) Total lung cellularity at 5 and 10 dpi. Data for 10 dpi were pooled from four independent experiments with 2–5 mice per group. Xist WT and Xist TcKO groups were compared using two-tailed Mann–Whitney tests. Data are shown as mean ± SD (**G–O**) Flow-cytometric analysis of lung immune and T cell populations at 10 dpi, showing frequency and cells/mg of lung for CD45^+^ immune cells (**G**), total CD3^+^ T cells (**H**), Foxp3^+^ Treg cells (**I**), total CD4^+^ T cells (**J**), CD4^+^ Naive T cells (**K**), CD4^+^ AgExp T cells (**L**), total CD8^+^ T cells (**M**), CD8^+^ Naive T cells (**N**), and CD8^+^ AgExp T cells (**O**). Each data point represents one mouse, and Xist WT and Xist TcKO groups were compared using two-tailed Mann–Whitney tests. Data are shown as mean ± SD. \**P* < 0.05, \*\**P* < 0.01; ns, not significant.

We infected female Xist TcKO and Xist WT mice, either on BALB/c or C57BL/6 backgrounds, aged 13-16 weeks with sublethal doses of influenza PR8 virus. Whole-lung viral titers at 3 dpi were comparable between Xist WT and TcKO mice, indicating that early viral burden was not significantly altered by Xist deletion in T cells (**Fig. 4B**). Surprisingly, we observed that female Xist TcKO mice on either background lost significantly less body weight at 7-8 dpi compared to WT controls (**Fig. 4C**). At 10 dpi, Xist WT lungs displayed prominent peri-bronchial and parenchymal inflammatory infiltrates, with substantial loss of normal alveolar airspace surrounding the conducting airways (**Fig. 4D; left**). In contrast, Xist TcKO lungs showed qualitatively reduced inflammatory infiltration and preservation of alveolar architecture (**Fig. 4D; right**). Consistent with the reduced pulmonary inflammation observed histologically, lung weights were significantly lower in Xist TcKO mice compared with WT mice at 10 dpi but not 5 dpi (**Fig. 4E**). However, lung cellularity normalized to tissue weight was comparable between genotypes at 5 and 10 dpi (**Fig. 4F**), suggesting that the reduced total lung cellularity in Xist TcKO mice primarily reflected decreased lung mass or fluid rather than a difference in cellular density.

To determine whether the attenuated pulmonary inflammation observed in Xist TcKO mice was associated with altered immune cell accumulation, we characterized lung immune cell populations by flow cytometry at 10 dpi. Despite reduced lung weights and inflammatory infiltrates in Xist TcKO mice, the frequency of total CD45^+^ immune cells and the number of CD45^+^ cells per mg of lung tissue were comparable between genotypes (**Fig. 4G**). Similarly, total CD3^+^ T cell frequencies and normalized cell numbers were unchanged (**Fig. 4H**). Further analysis revealed no significant differences in the frequencies or normalized numbers of Foxp3^+^ regulatory T cells, total CD4^+^ T cells, or CD4^+^ Naive and AgExp T-cell subsets (**Fig. 4I–L**). In contrast, Xist TcKO mice exhibited a significantly reduced frequency of total CD8^+^ T cells among CD3^+^ T cells, accompanied by an increased frequency of CD8^+^ Naive T cells. However, neither total CD8^+^ T cell numbers per mg of lung tissue nor the frequency and normalized abundance of CD8^+^ AgExp T cells differed significantly between genotypes (**Fig. 4M–O**). Lung CD19^+^ B cell frequency and total numbers were similar between WT and Xist TcKO mice (**Fig. S5D**), and there were comparable levels of influenza-specific HAI antibodies at 30 dpi (**Fig. S5E**). Examination of spleen tissue from infected female mice at 10 dpi revealed increased spleen masses for Xist TcKO mice but no differences with cellularity (**Fig. S5F**). Splenic T cell compartments were similar between infected WT and Xist TcKO animals (**Fig. S5G-M**). Together, these findings indicate that T cell specific *Xist* deletion is associated with changes in pulmonary CD8^+^ T cell composition at 10 dpi, despite largely preserved immune cell density and CD4^+^ T cell populations.

### Lung T cells from Xist TcKO females have increased expression of some XCI escape genes and antiviral autosomal genes

Next, we examined how Xist deletion affected gene expression in 10 dpi T cell subsets from influenza-infected WT and Xist TcKO lungs T cells using bulk RNA sequencing (**Fig. 5A**). Because XCI is random, we are unable to perform allele-specific transcriptional profiling in Xist TcKO mice, similar to *Xist* deletion in B cells^14,18,25^. While *Xist* deletion did not alter X-linked or autosomal differential gene expression in CD4^+^ Naive T cells (**Fig. 5B-C**), we found that there were significant differentially expressed X-linked and autosomal genes for CD8^+^ Naive and CD4^+^ and CD8^+^ AgExp T cells (**Fig. 5D-I**). Looking specifically at X-linked genes using a 1.5-fold cutoff, we found ∼5 X-linked genes significantly upregulated in AgExp T cells and CD8^+^ Naive T cells, notably the flu-specific XCI escape genes *Tktl1* and *Cxcr3* in AgExp T cells (**Fig. 5D, 5F, 5H**). Notably, three of the five X-linked genes upregulated in Xist TcKO T cells are T cell XCI escape genes^15^.

**Figure 5.**
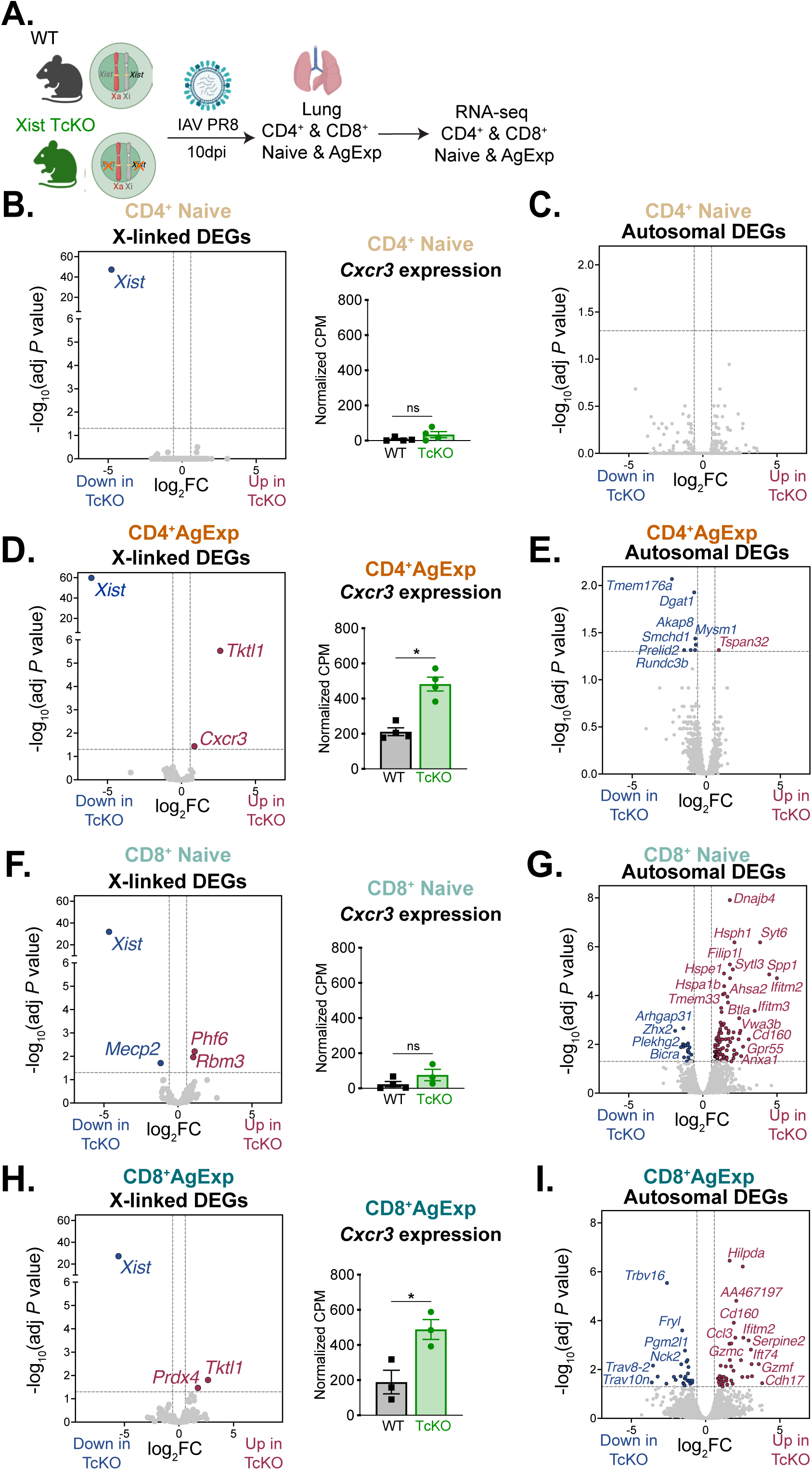
T cell-specific *Xist* deletion alters transcriptional profiles for X-linked and autosomal genes in pulmonary T cell subsets following influenza infection. (**A**) Experimental design for transcriptional profiling of flu infected Xist TcKO and WT female mice. Lung CD4^+^ and CD8^+^ Naive and AgExp T cells were isolated at 10 dpi for bulk RNA-seq. (**B–I**) Volcano plots showing X-linked (B,D,F,H) and autosomal differentially expressed genes (DEGs) (C,E,G,I) between Xist TcKO and WT cells in CD4^+^ Naive (B,C), CD4^+^ AgExp (D,E), CD8^+^ Naive (F,G), and CD8^+^ AgExp (H,I) T-cell populations. Differential expression was analyzed using DESeq2 with the contrast TcKO vs. WT. DEGs were defined by adjusted *P* ≤ 0.05 and absolute log2 fold-change ≥ log2(1.5). Red indicates genes significantly upregulated in TcKO cells, blue indicates genes significantly downregulated in TcKO cells, and gray indicates genes not meeting the differential-expression criteria. *Cxcr3* expression (shown as normalized CPM) in the indicated CD4^+^ (B,D) and CD8^+^ Naive and AgExp (F,H) T-cell populations. Each data point represents one mouse (n = 4/group). WT and TcKO groups were compared using two-tailed Mann–Whitney tests. Data are shown as mean ± SEM. \**P* < 0.05; ns, not significant.

*Xist* deletion resulted in relatively few autosomal transcriptional changes in CD4^+^ Naive and AgExp T cells, whereas both CD8^+^ T cell subsets exhibited substantially greater alterations in autosomal gene expression (**Fig. 5C, 5E, 5G, 5I**). Notably, CD8^+^ Naive T cells displayed increased expression of interferon-stimulated genes (*Ifitm2*, *Ifitm3*, and *Mx1*), cellular stress responses, and protein homeostasis, including *Dnajb4, Hsph1, Hspa1b*, and *Hspe1* (**Fig. 5G**). CD8^+^ AgExp T cells exhibited increased expression of genes involved in antiviral responses (*Ifitm2, Gzmc, Gzmf*), T cell activation and immune regulation (*Cd160*), inflammatory signaling (*Ccl3*), and hypoxia-associated metabolism (*Hilpda*). These findings reveal subset-specific transcriptional consequences of *Xist* deletion, with particularly pronounced alterations in CD8^+^ T cells. The enrichment of antiviral, metabolic, and cellular stress-response genes raises the possibility that *Xist* deletion modifies CD8^+^ T-cell adaptation and function during influenza infection.

### AgExp T cells from infected Xist TcKO females have increased biallelic expression of Cxcr3 mRNA and increased levels of Cxcr3 protein

Because AgExp T cells from influenza-infected Xist TcKO mice exhibited significantly higher levels of *Cxcr3* transcripts, we investigated the allelic frequency of *Cxcr3* expression using single-molecule RNA FISH (smFISH). We designed primary *Cxcr3* exonic (42 unique probes) and intronic (21 unique) probes using the PaintSHOP probe design tool^26^, each containing a 32 bp sequence for binding specific fluorescent secondary probes. For detection of exonic *Cxcr3* transcripts, we used secondary probes conjugated to ATTO-647N, and for detection of intronic *Cxcr3* transcripts we used ATTO-565 conjugated secondary probes. We infected cohorts of WT and Xist TcKO mice with influenza, and sorted CD4^+^Cxcr3^+^ and CD8^+^Cxcr3^+^ AgExp T cell populations from lungs and spleen at 10 dpi for smFISH. We developed a MATLAB program for semi-automated quantification of monoallelic and biallelic Cxcr3 RNA puncta of T cell nuclei that was integrated as a custom macro within ImageJ. The MATLAB app consisted of 3 steps for tracking monoallelic and biallelic puncta: (1) selection of morphologically acceptable nuclei; (2) deduplication of nuclei from potentially overlapping fields; (3) quantification of allelic nascent transcription (distinguishing mature exonic vs nascent intronic transcription activities) events using maximum projection images. Imaging WT and Xist TcKO Cxcr3^+^ AgExp T cells from infected lungs and spleen using *Cxcr3*-specific probes, we detected both monoallelic and biallelic *Cxcr3* expression in AgExp T cells (**Fig. 6A,B and Fig. S6A,B**), supporting that *Cxcr3* escapes XCI in *in vivo* and *in vitro* activated T cells (**Fig. 3** and ^15^). Using the MATLAB app to quantify the presence of nascent transcription of monoallelic and biallelic Cxcr3 RNA signals from both exonic and intronic probes, we found that Xist TcKO CD4^+^ and CD8^+^ AgExp T cells from both lung and spleen had significantly higher frequencies of biallelic nuclei compared to WT cells (**Fig. 6C-F**). CD4^+^ and CD8^+^ AgExp T cells from infected Xist TcKO exhibited ∼40% biallelic *Cxcr3* expressing nuclei in lung (**Fig. S6C,D**) and ∼30-35% biallelic Cxcr3 expression in spleen (**Fig. S6E,F**) compared to 15-20% biallelic expression in WT lung and spleen.

**Figure 6.**
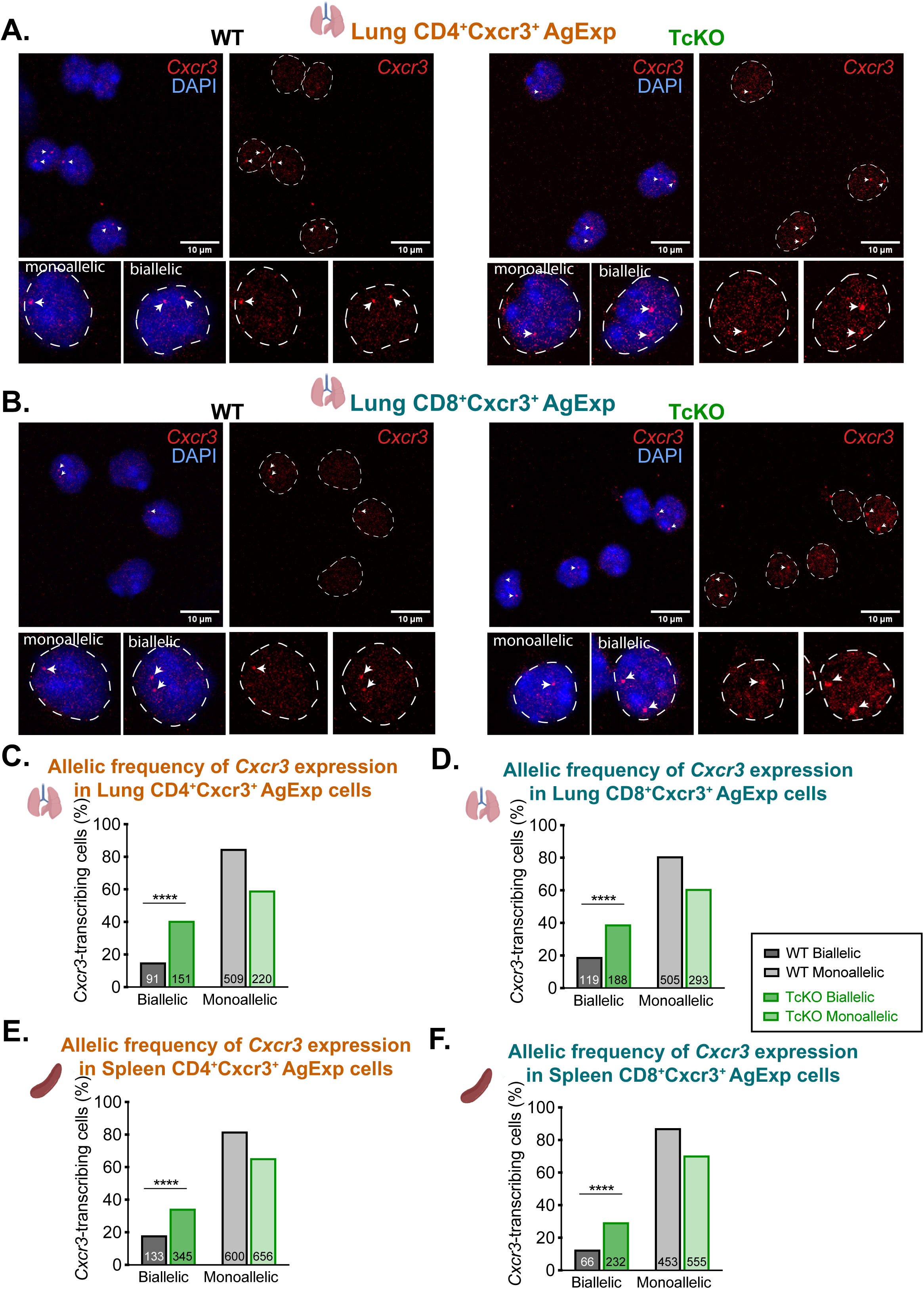
CD4^+^ and CD8^+^ AgExp T cells from lungs and spleens of infected Xist TcKO mice exhibit significantly increased biallelic *Cxcr3* expression. (**A–B**) Representative *Cxcr3* RNA smFISH images of lung CD4^+^ (A) and CD8^+^ (B) AgExp Cxcr3^+^ T cells from WT and Xist TcKO mice at 10 dpi, and signals for *Cxcr3* intron-specific probes (Cy3 channel only) are shown. Representative examples of monoallelic and biallelic *Cxcr3* nascent transcription are shown below. White arrowheads indicate pinpoints of nascent *Cxcr3* transcription. Scale bars, 10 μm. Staining with exonic *Cxcr3* probes was performed (not shown) and used for MATLAB puncta quantification to call nascent pinpoint transcription. (**C–F**) Percentage of monoallelic and biallelic *Cxcr3*-transcribing nuclei for WT and Xist TcKO mice for CD4^+^ (C, E) and CD8^+^ (D, F) AgExp CXCR3^+^ T cells from lung (C–D) and spleen (E–F). Numbers within each bar indicate the total number of nuclei counted. Data represent two independent experiments with 3–4 mice for each genotype and T cell subset, except lung Xist TcKO (n=2 mice, comprising 1 mouse and 3 pooled mice). Spleen samples were analyzed using individual mice for each genotype. WT and TcKO distributions were compared using two-sided Chi-square tests. \*\*\*\**P* < 0.0001.

Next, we investigated whether T cell subsets from infected Xist TcKO mice, exhibiting biallelic *Cxcr3* expression, express more Cxcr3 protein. We performed multiparametric flow cytometry of lung and spleen CD4^+^ and CD8^+^ T cells at 10 dpi from Xist TcKO and WT mice. UMAP visualization identified distinct populations corresponding to Naive (CD44^-^ CD62L^+^), AgExp (CD44^+^CD62L^+^), and CD4^+^Foxp3^+^ Treg cells, which were present in both WT and Xist TcKO mice, and Cxcr3 protein was enriched within the CD4^+^ AgExp T-cell compartment. (**Fig. 7A**). Although the overall distribution of CD4^+^ T cell subsets was largely preserved, Xist TcKO mice exhibited increased frequencies and numbers of Cxcr3^+^ cells within both CD4^+^ Naive and AgExp T-cell subsets in the lung (**Fig. 7B,C**). In contrast, UMAP visualization of flow cytometric analysis of CD8^+^ T cells revealed an apparent shift in cellular distribution toward the Naive compartment in Xist TcKO mice, consistent with the increased frequency of CD8^+^ Naive T cells in lung observed by conventional gating (**Fig. 7D**, **Fig. 4N**). Notably, *Xist* deletion significantly increased the frequency of Cxcr3^+^ cells within the CD8^+^ AgExp T cell compartment, accompanied by increased Cxcr3 expression intensity, while the number of Cxcr3^+^ CD8^+^ AgExp T cells per mg of lung tissue was not significantly altered (**Fig. 7E–F**). To determine whether the increased Cxcr3 protein and Cxcr3^+^ T cells observed in Xist TcKO lungs extended beyond a local tissue response, we examined splenic CD4^+^ and CD8^+^ T-cell populations at 10 dpi. In contrast to the pronounced increases in Cxcr3^+^ T-cell frequencies observed in the lung, *Xist* deletion did not significantly alter the frequencies or numbers of Cxcr3^+^ CD4^+^ Naive or AgExp T cells in the spleen (**Fig. S7A-C**). Similarly, the frequency of Cxcr3^+^ CD8^+^ AgExp T cells was unchanged, although Xist TcKO mice exhibited a modest but significant increase in the frequency of Cxcr3^+^ CD8^+^ Naive T cells (**Fig.S7D-F**). Together, these findings indicate that *Xist* deletion modifies Cxcr3 mRNA and protein expression across pulmonary T-cell subsets, with particularly pronounced changes in CD4^+^ and CD8^+^ AgExp T cells.

**Figure 7.**
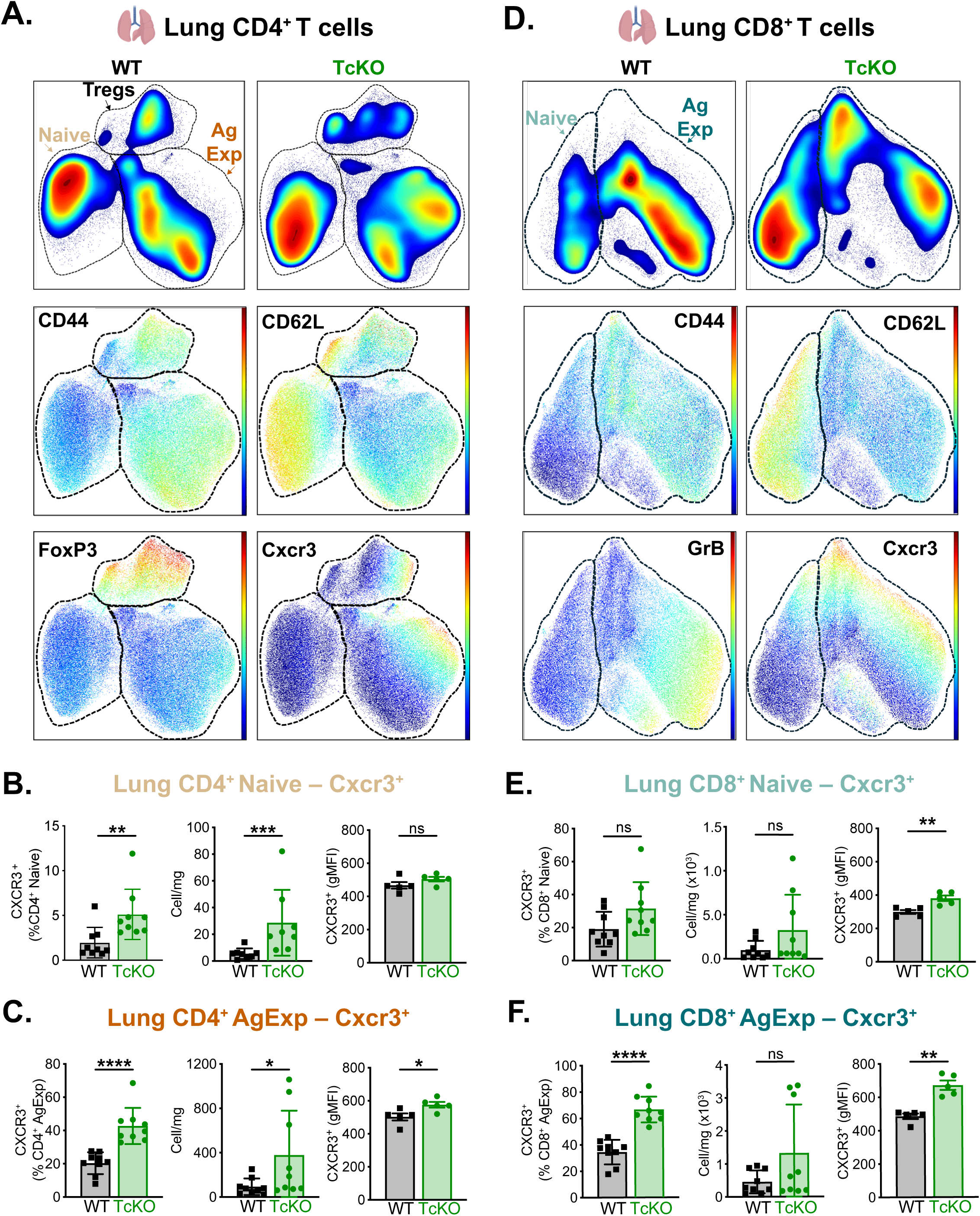
T cell-specific *Xist* deletion enhances Cxcr3 protein levels in lung T cell compartment following influenza infection. (**A,D**) Representative UMAP plots of lung CD4^+^ (A) and CD8^+^ (D) T cells from WT and Xist TcKO mice at 10 dpi. (A) Lung CD4^+^ Naive, AgExp, and Treg populations and the expression of the indicated markers, including Cxcr3. (**B,C**) Quantification of Cxcr3^+^ CD4^+^ Naive (B) and AgExp (C) T cells from flu-infected lungs of WT and Xist TcKO mice. The frequency and number of Cxcr3^+^ cells/mg of lung and Cxcr3 protein (as gMFI) are shown. (**E,F**) Frequency and number (cells/mg lung) of Cxcr3^+^ CD8^+^ Naive (E) and AgExp (F) T cells from flu-infected lungs of WT and Xist TcKO mice. Frequency and cell number data were pooled from two independent experiments with 4–5 mice/group and are shown as mean ± SD. Cxcr3 gMFI data are representative of two independent experiments with n = 5 mice per group and are shown as mean ± SEM where each data point represents one mouse. WT and Xist TcKO groups were compared using two-tailed Mann–Whitney tests. \**P* < 0.05, \*\**P* < 0.01, \*\*\**P* < 0.001, \*\*\*\**P* < 0.0001; ns, not significant.

### Xist deletion enhances IFN-γ and TNF production by pulmonary and splenic AgExp T cells from influenza-infected mice

Next, we investigated whether the increased Cxcr3 protein levels observed in *Xist*-deficient pulmonary T cells was associated with enhanced effector function, particularly IFN-γ and TNF cytokine production following *ex vivo* stimulation. Lung cells isolated from influenza-infected Xist WT and TcKO mice at 10 dpi were stimulated with PMA and ionomycin or cultured in media alone for 4 h, followed by intracellular cytokine staining and flow cytometric analysis (**Fig. 8A**). UMAP visualization of gated CD4^+^ and CD8^+^ T cells revealed heterogeneity among Naive, AgExp, and Treg T-cell populations, with differential expression of IFN-γ, TNF, Cxcr3, CD40L, and granzyme B (GrB) (**Fig. 8B,C**). Within the CD4^+^ Naive T-cell compartment, *Xist* deletion increased the frequency of Cxcr3^+^IFN-γ^+^ cells, whereas the frequencies of total IFN-γ^+^, Cxcr3^+^, and IFN-γ^+^TNF^+^ polyfunctional cells were not significantly altered (**Fig. 8D**). In contrast, CD4^+^ AgExp T cells from Xist TcKO mice exhibited significantly increased frequencies of IFN-γ^+^, Cxcr3^+^, Cxcr3^+^IFN-γ^+^, and IFN-γ^+^TNF^+^ polyfunctional cells (**Fig. 8E**). Similar changes were observed in CD8^+^ T cells, with increased frequencies of IFN-γ^+^ and Cxcr3^+^IFN-γ^+^ cells in the Naive compartment (**Fig. 8F**). Notably, CD8^+^ AgExp T cells exhibited significant increases in all four populations, including IFN-γ^+^TNF^+^ polyfunctional cells (**Fig. 8G**). Together with the increased *Cxcr3* transcript abundance (**Fig. 6**) and Cxcr3 protein expression observed in unstimulated pulmonary T cells (**Fig. 7**), these findings indicate that *Xist* deletion is associated with enhanced Type 1 cytokine-producing capacity in CD4^+^ and CD8^+^ AgExp T cells during influenza infection.

**Figure 8.**
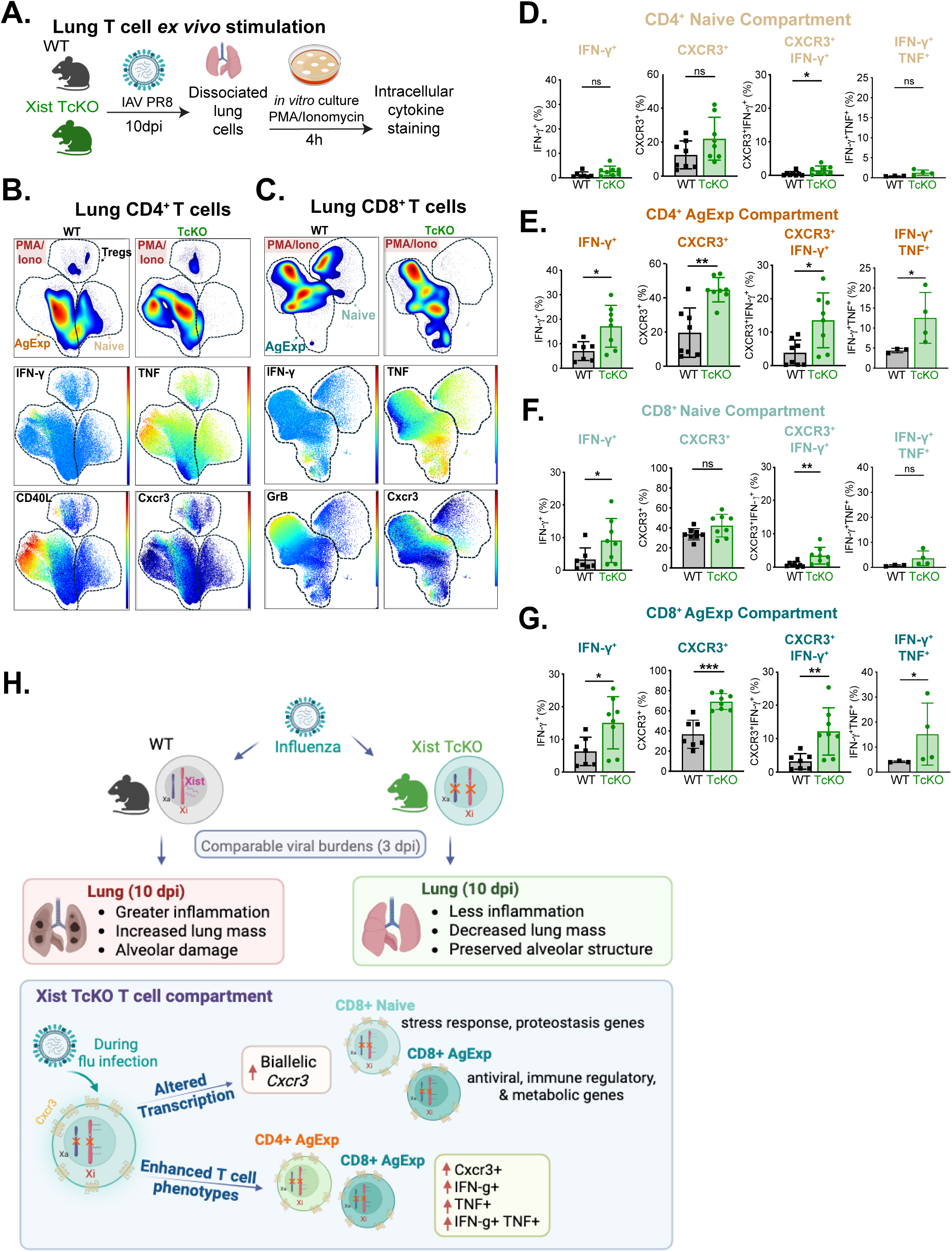
*Xist* deletion enhances the cytokine production potential of AgExp T cells from infected lungs. **(A)** Experimental design for *ex vivo* T-cell stimulation. Female Xist WT and Xist TcKO mice were infected with influenza PR8, and dissociated lung cells were stimulated with PMA/Ionomycin for 4h at 10 dpi, followed by intracellular cytokine staining. (**B, C**) Representative UMAP plots of lung CD4^+^ (B) and CD8^+^ (C) T cells from WT and Xist TcKO mice following *ex vivo* stimulation, showing Naive, AgExp, and Treg populations (for CD4^+^ cells). Expression levels of IFN-γ, TNF, CD40L, and GrB are shown overlaid on UMAP plots. (**D-G**) Quantification of frequencies of CD4^+^ Naive (D), CD4^+^ AgExp (E), CD8^+^ Naive (F), and CD8^+^ AgExp (G) for IFN-γ^+^, Cxcr3^+^, IFN-γ^+^Cxcr3^+^, and IFN-γ^+^TNF^+^ cells following *ex vivo* stimulation. Data were pooled from two independent experiments with 4–5 mice per group per experiment. Each data point represents one mouse. WT and TcKO groups in (D–Q) were compared using two-tailed Mann–Whitney tests. Data are shown as mean ± SD. \**P* < 0.05, \*\**P* < 0.01; ns, not significant. **(H)** Model. Sublethal influenza infection of female WT and Xist TcKO mice, despite comparable viral burdens at 3 dpi, results in greater lung injury for WT animals at 10 dpi. Within the Xist TcKO compartment, during flu infection there is more biallelic *Cxcr3* expression and elevated protein levels, and CD8^+^ T cells exhibit increased expression of antiviral, immune regulatory, stress response, and metabolic genes. Xist TcKO CD4^+^ and CD8^+^ AgExp have greater potential to produce inflammatory cytokines IFN-γ and TNF. Thus, a T cell-specific deletion of *Xist* reduces influenza-associated pulmonary pathology while enhancing antigen-experienced T cell cytokine producing potential.

To determine whether the enhanced cytokine-producing capacity of *Xist*-deficient T cells extended beyond the site of infection, we performed similar *ex vivo* stimulation experiments using splenocytes isolated from influenza-infected WT and Xist TcKO mice at 10 dpi (**Fig. S8A**). Following PMA/ionomycin stimulation, UMAP visualization revealed phenotypically distinct CD4^+^ Naive, AgExp, and Treg populations (**Fig. S8B**), as well as CD8^+^ Naive and AgExp T-cell populations (**Fig. S8C**). Within the splenic CD4^+^ and CD8^+^ Naive T-cell compartments, the frequencies of Cxcr3^+^, Cxcr3^+^IFN-γ^+^, and IFN-γ^+^TNF^+^ polyfunctional cells were comparable between genotypes (**Fig. S8B,C**). However, Xist TcKO mice exhibited significantly increased frequencies of Cxcr3^+^, Cxcr3^+^IFN-γ^+^, and IFN-γ^+^TNF^+^ polyfunctional cells within both splenic CD4^+^ and CD8^+^ AgExp T-cell populations (**Fig. S8D,E**). To determine whether enhanced cytokine-producing capacity was accompanied by increased degranulation potential, we examined CD107a surface expression in pulmonary T cells following *ex vivo* stimulation. Neither the frequency of CD107a^+^ cells nor the proportion of Cxcr3^+^CD107a^+^ cells differed significantly between WT and Xist TcKO mice within CD4^+^ or CD8^+^ Naive and AgExp T-cell subsets (**Fig. S8F,G**). Thus the enhanced cytokine-producing capacity associated with *Xist* deletion was not accompanied by a detectable increase in CD107a surface expression under the stimulation conditions examined. Collectively, these results indicate *Xist* deletion enhances cytokine-producing potential in both pulmonary and splenic AgExp T cells following influenza infection.

## DISCUSSION

Here, we investigated how influenza infection impacts the transcriptional and epigenomic profile of the Xi in Naive and AgExp T cells, and how *Xist* deletion influences T cell responses to influenza infection. Sequential RNA FISH/IF, together with allele-specific transcriptional and epigenomic profiling, revealed that the Xi in pulmonary T cell subsets exhibits infection-specific XCI escape and variable enrichment of H3K27me3 and H2AK119Ub modifications. Using a novel T cell-specific *Xist* deletion mouse model, we found that Xist TcKO mice infected with influenza exhibit less lung injury despite comparable viral titers, increased biallelic *Cxcr3* expression and Cxcr3 protein levels, and enhanced Cxcr3-associated IFN-γ/TNF responses (**Fig. 8H**). Our study describes a novel role for Xist RNA in T cells in regulating cytokine production, thereby influencing pulmonary responses to influenza infection.

We found that flu infection results in much less cytological enrichment of Xist RNA and H2AK119Ub foci at the Xi in CD4^+^ and CD8^+^ T cells across lung and peripheral tissues at both 5 and 10 dpi, as compared to what we previously observed for *in vitro* CD3^+^ T cell stimulation^15,16^. The majority of Xist RNA signals resembled pinpoint clusters rather than canonical ‘cloud’-like structures in T cell subsets from flu-infected mice, while *in vitro*-stimulated T cells at 48-72 hrs post-stimulation result in higher frequencies of nuclei with larger clouds that encompass the nuclear area of the Xi^17^. We observed more Xist RNA signal in sites and times of higher antigen availability, specifically in the dLN at both timepoints and in the lung at 5 dpi, suggesting that T cell receptor stimulation is also the critical signal driving enrichment at the Xi *in vivo*. Using allele-specific CUT&RUN profiling, we observed that the Xi has low levels of H2AK119Ub enrichment across Naive and AgExp T cell compartments, at levels resembling the Xa following infection. Yet the Xi has ∼2-3x higher enrichment of H3K27me3 compared to Xa across T cell subsets from infected female mice, supporting a functional role for H3K27me3 marks as imprints for transcriptional silencing on the Xi, independent of Xist RNA association with the Xi, in lymphocytes^18^. We have previously proposed that DNA methylation (DNAm) at CpG promoters, together with H3K27me3 marks, are required to maintain transcriptional silencing of Xi-repressed genes in B cells, which also exhibit dynamic XCI maintenance mechanisms^18^. It is likely that Xi promoter regions in pulmonary T cell subsets from infected mice will exhibit high levels of DNAm and that infection will not significantly alter DNAm at repressed genes. Additional work is needed to investigate whether DNAm levels at Xi promoters are stable following influenza infection and determine if reducing DNAm across Xi will influence Xi gene reactivation.

We compared enrichment changes for H2AK119Ub and H3K27me3 at Xi genes between AgExp and Naive T cells at 10 dpi, corresponding to the peak of T effector responses after influenza infection. We were surprised that ∼200-225 Xi genes lost H3K27me3 marks in AgExp compared to Naive T cells, while ∼67-149 Xi genes gained H2AK119Ub modifications. It is possible that reduced H3K27me3 modifications at these Xi genes alters the heterochromatic environment at these regions, resulting in more euchromatic chromatin that is permissive for Xi reactivation. Our allele-specific transcriptional profiling revealed that ∼50-63 XCI escape genes are expressed from the Xi across pulmonary T cell subsets in infected female mice. The majority of XCI escape genes are expressed in both Naive and AgExp T cells, indicating that this increased dosage is important for T cell function irrespective of activation status. As XCI escape genes are usually expressed at higher levels in female compared to male cells, it is likely that these XCI escape genes contribute to observed sex differences with T cell-mediated responses to influenza infections. In support, the constitutive XCI escape gene *Kdm6a* is expressed at higher levels in female CD4^+^ T cells and NK cells and contributes to sex differences in susceptibility to autoimmune disease and antiviral responses^27,28^. We found that *Cxcr3* is biallelically expressed in AgExp CD4+ and CD8+ T cells from flu-infected lungs, yet intriguingly gains genic H3K27me3 and H2AK119Ub as compared to Naïve CD4+ and CD8+ T cells. Using a *Leishmania mexicana* model of infection and a dual fluorescent Cxcr3 reporter system, female biallelic *Cxcr3*-expressing T cells have enhanced inflammatory characteristics^20^. Our study confirms that Xi-specific *Cxcr3* expression is a feature of T cell immune responses to infection, and suggests that activation-dependent expression may be epigenetically controlled by a bivalent-like chromatin state at some XCI escape genes, warranting further investigation.

One of the most striking findings of our study is that a T cell-specific deletion of *Xist* protects female mice from influenza-induced weight loss and pulmonary injury. Despite comparable viral burdens at 3 dpi, Xist TcKO mice exhibited significantly reduced weight loss, diminished pulmonary inflammatory infiltrates, and preservation of alveolar architecture at 10 dpi, the late effector phase of infection. Infected Xist TcKO mice also reduced lung weights, consistent with attenuated pulmonary inflammation and tissue injury. Surprisingly, the protective phenotype was not associated with substantial changes in pulmonary immune cell composition or abundance, as the frequencies and numbers of total CD45^+^ immune cells, CD3^+^ T cells, and CD4^+^ and CD8^+^ AgExp T cells normalized to lung tissue weight were largely unchanged. However, Xist TcKO mice exhibited a reduced frequency of total CD8^+^ T cells and an increased proportion of CD8^+^ Naïve T cells, suggesting altered composition of the pulmonary T cell compartment. Because influenza-associated lung injury results from both direct viral damage and host inflammatory responses, these findings raise the possibility that Xist deficiency modifies the nature of the pulmonary immune response rather than simply limiting T cell recruitment or accumulation. Thus, Xist-dependent regulation of T cell responses may contribute to the severity of influenza-associated pulmonary pathology independently of major changes in local T cell abundance.

Unexpectedly, the reduced pulmonary pathology in Xist TcKO mice was accompanied by enhanced cytokine-producing potential in CD4^+^ and CD8^+^ AgExp T cells. Following *ex vivo* stimulation, Xist-deficient pulmonary T cells exhibited increased frequencies of IFN-γ and IFN-γ/TNF-producing AgExp T cells, consistent with enhanced effector cytokine responses. These functional changes were associated with increased biallelic *Cxcr3* transcription and elevated pulmonary Cxcr3 protein expression, suggesting that altered X-linked gene dosage may contribute to the enhanced functional state of *Xist*-deficient T cells. Notably, increased cytokine-producing potential was also observed in splenic AgExp T cells, indicating that the consequences of *Xist* deletion extend beyond the site of infection in the lung. Despite these enhanced cytokine responses, Xist TcKO mice exhibited reduced lung inflammation and tissue injury, challenging the assumption that greater T cell effector potential necessarily exacerbates immunopathology. Rather, our findings suggest that T cell activation and cytokine-producing capacity can be uncoupled from the severity of pulmonary tissue damage. Whether this protection reflects altered T cell localization, effector T cell differentiation, or regulation of inflammatory responses within the infected lung remains to be determined. Together, these findings identify *Xist* as an unexpected regulator of T cell function during viral infection and suggest that *Xist*-dependent modulation of X-linked gene expression influences the balance between antiviral effector responses and pulmonary immunopathology.

## METHODS

### Mice

Female wildtype C57BL/6J and BALB/c mice were used for imaging of Xist RNA and heterochromatic marks in T cell subsets post PR8 influenza infection. Female F1 *Mus musculus castaneus* X C57BL/6J *Xist^fl/+^* (F1 mus x cast) mice were used for all allele-specific CUT&RUN and RNA-seq experiments, and were generated by mating Xist^fl/fl^ female mice to male b-actin Cre (B6N.FVB-*Tmem163^Tg(ACTB-cre)2Mrt^*/CjDswJ; strain# 019099, Jackson), then Xist^fl/+^ female mice were mated to male *Mus musculus castaneus* (Cast; strain# 000928, Jackson) mice to generate female F1 mus x cast mice (Sierra *et al.* ^14^). Female Xist cKO mice with a T cell specific deletion were used for *in vivo* Xist deletion experiments, and were generated by mating Xist^fl/fl^ mice on a C57BL/6 or BALB/c background to CD4-cre mice (B6.Cg-Tg(Cd4-cre)1Cwi/BfluJ; Jackson Laboratory Strain 022071; we backcrossed this Cd4-cre strain onto BALB/c background for 12 generations. Xist+/+ and CD4-Cre^+/-^ were used as controls for these experiments. Mice were maintained at the Penn Vet animal facility, and experiments were approved by the University of Pennsylvania Institutional Animal Care and Use Committee (IACUC).

### Viral stocks and infections

Influenza virus A/PR8 stocks were kindly provided by L.C. Eisenlohr and M.Hogan (1.64×10^6 PFU/mL). Sublethal doses of virus (3-10 PFU) were administered intranasally under light isoflurane anesthesia in 30 μL of PBS. Mice were 20.4 ± 5.0 weeks old at the time of infection. All infected mice were monitored daily for weight loss, hunched posture, ruffled fur, and lack of movement and euthanized when humane endpoints were reached.

### Tissue collection, processing, & histopathology

Following euthanasia by carbon dioxide inhalation, lungs were perfused with 1x PBS, weighed, minced with a razor blade, and digested under rotation for 1 hour at 37°C in RPMI 1640 medium (Invitrogen) containing Liberase TL (0.1 WU/mL; Roche, 274113) and DNase I (30 µg/mL; Roche, 10104159001). The digested tissue was passed through a 70-µm basket filter, red blood cells were lysed using ACK Lysis Buffer (Quality Biological, 118-156-721), and cells were resuspended in FACS buffer (0.5% BSA in PBS). Spleens and mediastinal lymph nodes were mechanically dissociated using glass slides, filtered through a 70-µm filter, RBC-lysed, and resuspended in FACS buffer. Blood collected by cardiac puncture was centrifuged at 10,000 x g to obtain serum, which was stored at −80°C. For tissue sectioning for histopathology, perfused whole lungs were inflated with 10% neutral buffered formalin (NBF; Sigma-Aldrich, HT501128-4L), fixed for 72 hours, and subsequently stored in 70% ethanol. Lungs were paraffin-embedded, sectioned, and stained with hematoxylin and eosin (H&E) by the PennVet Comparative Pathology Core. Slides were imaged at 20x magnification using an Aperio VERSA 200 slide scanner.

### T cell subset isolation, fluorescence-activated cell sorting (FACS), and *in vitro* stimulation

Single-cell suspensions were stained with LIVE/DEAD Fixable Aqua or Near-IR (Life Technologies, L34965 or L34992) and Fc receptors were blocked with anti-CD16/CD32 (clone 2.4G2; BD Biosciences, 553142). For imaging and sequencing experiments, CD4⁺ and CD8⁺ naive and antigen-experienced (AgExp) T cells were stained in FACS buffer using antibodies against CD3ε (17A2; BV785, 100231, or PE-Dazzle594, 100245), CD4 (GK1.5; PE-Cy7, 100422, or AF647, 100424), CD8α (53-6.7; BV421, 100737, or PerCP-Cy5.5, 100734), CD44 (IM7; AF700, 103026, or PE, 103024), and CD11a (I21/7; AF488, 153106), all from BioLegend. Following exclusion of dead cells and gating on singlets, CD4⁺ and CD8⁺ T cells were identified based on CD3ε, CD4, and CD8α expression. Naive T cells were defined as CD44⁻CD11a⁻, and AgExp T cells as CD44⁺CD11a⁺. For *Cxcr3* single-molecule RNA FISH (smFISH) experiments, T cells from the lung and spleen were enriched by magnetic-activated cell sorting (MACS) using the Pan T Cell Isolation Kit II (Miltenyi Biotec, 130-095-130), according to the manufacturer’s instructions. Enriched cells were stained as above, with anti-CXCR3 (CXCR3-173; PE-Dazzle594; BioLegend, 126533). CD4⁺ and CD8⁺ AgExp T cells were subsequently sorted by FACS into CXCR3⁺ and CXCR3⁻ populations. All cell sorting was performed using a BD FACSAria Fusion cell sorter.

For confirmation of Xist RNA cKO in T cells by Xist RNA FISH and heterochromatic mark immunofluorescence, murine splenic CD3⁺ T cells were isolated from WT and Xist^fl/fl^;CD4-cre^+^ mice using CD3⁺ T cell enrichment columns (R&D Systems, MTCC25). T cells were cultured at 37°C in RPMI 1640 (Invitrogen) containing 10% fetal bovine serum (FBS; Gemini), 0.1% β-mercaptoethanol (Invitrogen), 1% nonessential amino acids (Invitrogen), 1% sodium pyruvate (Invitrogen), and 1% penicillin-streptomycin (Invitrogen) and were stimulated with plate bound αCD3 (1 μg/ml; Bio X cell, BE0001–1-5MG) and soluble αCD28 (10 μg/ml; Bio X cell, clone 37.51) for 72 hours.

### Flow cytometry and intracellular staining

Single-cell suspensions were stained and Fc-blocked as described above. For surface immunophenotyping, cells were stained in FACS buffer using antibodies against CD4 (GK1.5; PE-Cy7, 100422), CD8α (53-6.7; PerCP-Cy5.5, 100734), CD44 (IM7; PE, 103024), CD3ε (145-2C11; FITC, 100305), CD19 (eBio1D3; APC-eFluor 780; eBioscience, 47-0193-82), and CD45 (30-F11; AF700, 103128), with all antibodies from BioLegend unless otherwise indicated. To assess cytokine production, degranulation, and activation markers in T cells, splenocytes or dissociated lung cells were analyzed either directly *ex vivo* or following *in vitro* stimulation with PMA (50 ng/mL) and ionomycin (500 ng/mL) or cultured with media alone for 4 hours in the presence of GolgiPlug and GolgiStop (BD Biosciences, 555029 and 554724, respectively), according to the manufacturer’s instructions. Cells were stained using different combinations of antibodies against IFN-γ (XMG1.2; BV421; BioLegend, 505829), CXCR3 (CXCR3-173; BV605; BioLegend, 126523), CD107a (1D4B; BV711; eBioscience, 407-1071-82), CD107b (ABL-93; FITC; eBioscience, 11-1072-81), CD45 (30-F11; BV785; BioLegend, 103149), CD11a (I21/7; AF488, 153106, or PE-Cy7, 153107; BioLegend), CD8α (53-6.7; PerCP-Cy5.5; BioLegend, 100734), CD40L/CD154 (MR1; PE-eFluor 610; eBioscience, 61-1541-80), CD4 (H129.19; PE-Cy5; BioLegend, 130312), CD3ε (17A2; PE-Fire 700; BioLegend, 100271), Foxp3 (150D; AF647; BioLegend, 320013, or FJK-16s; PE; eBioscience, 12-5773-80), CD44 (IM7; AF700; BioLegend, 103026), CD62L (MEL-14; BV650; BioLegend, 104453), granzyme B (GB11; AF647; BioLegend, 515405), TNF (MP6-XT22; APC-Cy7; BioLegend, 506343), and CD19 (eBio1D3; APC-eFluor 780; eBioscience, 47-0193-82). For intracellular staining, cells were fixed and permeabilized using the eBioscience Foxp3/Transcription Factor Staining Buffer Set (00-5523-00) according to the manufacturer’s instructions. CD107a/b were added during stimulation. Samples were acquired using a BD LSRFortessa flow cytometer and analyzed using FlowJo v10.10. Uniform Manifold Approximation and Projection (UMAP) projections were generated using OMIQ (Dotmatics). FCS files from WT and TcKO lung and spleen samples (n = 4 mice/group) were imported into OMIQ. Total CD4⁺ and CD8⁺ T cells were gated and analyzed separately. For a given T cell subset, WT and TcKO samples were combined into a single UMAP embedding, which was derived using markers (CD107b, IFN-γ, CXCR3, CD62L, CD107a, GrB, CD44, TNF, FoxP3, CD40L, and CD11a) and an equal number of cells per biological sample, as specified in the corresponding figure legend. UMAP projections were generated using 15 neighbors, with a minimum distance of 0.4, using a Euclidean distance metric. Cell distributions were visualized as filled contour density plots. The fluorescence intensities of each marker were also projected of each UMAP embedding. Naive, AgExp, and Treg T-cell populations were identified based on CD44, CD62L, and Foxp3 expression patterns

### Xist RNA fluorescent in situ hybridization (FISH) and Immunofluorescence (IF)

Sequential *Xist* RNA FISH and H2AK119Ub IF were performed as previously described^22^. Briefly, Xist RNA FISH was performed using Cy3-labeled oligonucleotide probes for Xist RNA^22,23^, and nuclei were counterstained using Vectashield with DAPI (Vector Lab H-1200). Sequential immunofluorescence for histone tail modifications was performed using primary antibodies for H2AK119Ub (Cell Signaling 8240S) and secondary antibodies AF488-conjugated goat anti-rabbit IgG (Abcam ab150077-500UG). Samples were imaged using Nikon Eclipse Microscope and patterns of Xist RNA and histone focal enrichment were quantified manually as previously described^15^.

### CUT&RUN

CUT&RUN for H3K27me3 and H2AK119Ub was performed on sorted CD4^+^ and CD8^+^ Naive and AgExp T cells from the lung of influenza infected F1 mus x cast female mice at 5 and 10 dpi using previously published methods^29^. In brief, 10uL/reaction of concanavalin A-coated beads (Cat #86057-3, Polysciences) were equilibrated with Binding Buffer (20mM HEPES, 10mM KCl, 1mM CaCl_2_, 1mM MnCl_2_) then resuspended in Wash Buffer (20mM HEPES, 150mM NaCl, 0.5mM Spermidine, 1x PIC (Cat #11697498001, Roche)). Approximately 10,000-30,000 sorted T cell were immobilized on beads in Wash Buffer by rotating for 1 hour at room temperature. Beads were resuspended in Antibody Buffer (20mM HEPES, 150mM NaCl, 0.5mM Spermidine, 1x PIC, 0.05% Digitonin, 2mM EDTA). Primary antibody for H3K27me3 (3 µg; Cat #39055, Active Motif), H2AK119Ub (∼1.5 µg; Cat # 8240S, Cell Signaling) or IgG (3 µg; Cat #A01008, GenScript) were added and beads were rotated at 4°C for 5 hr. Beads were washed with and resuspended in Dig-Wash Buffer (20mM HEPES, 150mM NaCl, 0.5mM Spermidine, 1x PIC, 0.05% Digitonin). 1200ng/mL of pA-MNase fusion protein (gift from Dr. Henikoff and Dr. K. Sarma) was added and beads were rotated at 4°C for 1 hr. Beads were washed twice then resuspended in Dig-Wash Buffer. Targeted digestion was performed at 0°C for 30 min by adding CaCl_2_ to 2mM. The digestion was stopped by adding one volume of 2X STOP Buffer (340mM NaCl, 20mM EDTA, 4mM EGTA, 0.02% Digitonin, 50ug/mL RNase A (Cat #3335399001, Roche), 50ug/mL Linear Acrylamide (Cat #K548, VWR), 2pg/mL Yeast Spike-In DNA). The target chromatin was released into the supernatant by shaking beads at 300rpm for 10 min at 37°C. The supernatant was then transferred to fresh tubes and incubated at 70°C for 10 min with SDS to 0.1% and 50ug of Proteinase K (Cat # EO0491, Thermo Scientific). DNA was then isolated and concentrations were measured using a Qubit system. For samples with insufficient concentration to obtain a Qubit reading, the entire sample was input into library preparation. For samples with a Qubit reading, 1.5ng was input into library preparation. Libraries were prepared using NEBNext Ultra II DNA Library Prep Kit for Illumina (Cat # E7645S, NEB) following vendor’s protocol and library quality was assessed using an Agilent TapeStation. CUT&RUN libraries were paired-end sequenced on a 300 cycle flow cell using a NextSeq2000 system.

### Data Processing for CUT&RUN

Adapters were removed using Trimmomatic (V0.36, https://github.com/usadellab/Trimmomatic). For allele-specific analyses, SNPsplit (0.5.0)^30^ was used to generate an “N-masked” version of the mouse reference genome (mm10), in which SNPs between the C57BL/6 and *Mus musculus castaneus* alleles are replaced with an N nucleotide in the mm10 genome. Reads were mapped to the “N-masked” genome using Bowtie2 (2.3.4.3) with the parameters [-q --local --very-sensitive-local --soft-clipped-unmapped-tlen --dovetail --no-mixed --no-discordant --phred33 -I 10 - X 1000]^31^. Duplicated reads were removed using Picard MarkDuplicates (2.17.11, https://broadinstitute.github.io/picard/) with option [REMOVE-DUPLICATES=true]. Reads mapped to regions from the ENCODE blacklist were removed. Reads overlapping SNPs were sorted into BL6 or Cast allele-specific BAM files were then generated using SNPsplit (0.5.0) with option [--paired]. CPM normalized bigwig files were generated from allele-specific and BAM files using deepTools bamCoverage (3.5.6) with options [--effectiveGenomeSize 2652783500 --normalizeUsing CPM --ignoreForNormalization chrX]^32^. CPM normalized biological replicates were combined with deepTools bigwigAverage (3.5.6) and visualized with Integrative Genomics Viewer (IGV, v2.14.0)^33^.

### Computational Analyses for CUT&RUN

#### Changes in Xi Heterochromatic Histone Mark Enrichment between groups

Visualization of heterochromatic mark enrichment between groups across the Xi was performed as previously described^34^. Reads from the *Cast* files (allele with the Xi) were counted in 10kb windows across the genome (generated with GenomicRanges (1.44.0)^35^) using featureCounts from the Rsubread package (2.6.4)^36^ with arguments [useMetaFeatures = F, isPairedEnd = T]. Consensus autosomal peaks were then use to calculate a normalization factor using the trimmed mean of M-values (TMM) method from the edgeR package (3.34.1)^37^. To determine consensus autosomal peaks, peaks were first called on Cast files using Sparse Enrichment Analysis for CUT&RUN (SEACR)^38^ with options [norm relaxed] and then merged across all timepoints and biological replicates for each mark using bedtools intersect (2.29.1)^39^ with parameters [-nonamecheck -a]. To correct for chromatin accessibility or map ability bias, 10kb windows with a large variability in the IgG samples (read counts exceeded ±2 standard deviations from the mean) were discarded from the analysis. Read counts for each 10kb window were then averaged between biological replicates. Delta enrichment was then calculated between timepoints within a subset (e.g. CD4^+^ Naïve 5 vs 10 dpi) or between activation states (e.g. CD4^+^ Naive vs CD4^+^ AgExp). Locally estimated scatterplot smoothing (LOESS) regression was performed on delta accumulation values across all 10kb bins.

#### d-score analysis

Genic regions were defined as 1kb upstream of the TSS (promoter region) to the TES. Intergenic regions were defined as 10kb windows (generated using GenomicRanges (1.44.0)^35^) that did not overlap gene bodies or their promoter regions (1kb upstream of TSS to TES). Reads were assigned to defined windows using featureCounts from the Rsubread package (2.6.4)^36^ with arguments [useMetaFeatures = F, isPairedEnd = T]. Regions with less than 5 allelic reads (reads^Cast^+ reads^BL6^) in any replicate were filtered out due to insufficient coverage. For each biological replicate, d-scores [(reads^BL6^/(reads^Cast^+ reads^BL6^))-0.5] were calculated for genic and intergenic regions. Biological replicates were then averaged for each timepoint and T cell subset. *p*-values were calculated using a Wilcoxon signed-rank test with Benjamini Hochberg correction.

#### Xi H2AK119Ub vs H3K27me3 enrichment across the Xi

Reads in 10kb bins on the *Cast* allele for H2AK119Ub and H3K27me3 were quantified and normalized using consensus autosomal peaks as described above. Read counts for each window were then averaged for each mark between biological replicates. LOESS regression across all 10kb bins was performed for each mark and plotted across the Xi.

#### Enrichment Analysis of H3K27me3 and H2AK119Ub at Xi genic regions

Cast reads in genic regions were quantified using featureCounts from the Rsubread package (2.6.4)^36^ with arguments [useMetaFeatures = F, isPairedEnd = T] and normalized using consensus autosomal peaks as described previously. Differentially enriched genes between Naive and AgExp cells were generated using the edgeR package^37^. To assess changes in enrichment at escape genes from each T cell subset (see below for escape gene identification), normalized H2AK119Ub and H3K27me3 reads at escape genes were filtered out and plotted on heatmaps.

### Allele-Specific and *Xist* TcKO RNA Sequencing

For allele-specific analysis of Xi gene expression post influenza infection, CD4^+^ and CD8^+^ Naive and AgExp T cells were sorted from lungs of F1 mus x cast mice at 10 dpi and stored in TRIzol (Thermo Fisher Scientific, 15596026). To determine the effect of Xist RNA deletion in T cells during infection, CD4^+^ and CD8^+^ Naive and AgExp T cells were sorted from lungs of BL6 WT and TcKO mice at 10 dpi and stored in TRIzol (Thermo Fisher Scientific, 15596026). RNA was isolated following the manufacturers protocol. F1 allele-specific libraries were prepared using the Illumina TruSeq Stranded Total RNA Stranded Ligation Ribo-Zero+ Kit (Illumina, 20040529) and run on an Illumina NextSeq 2000 sequencer (150-bp paired-end reads). TcKO libraries were prepared using the SMARTer Stranded Total RNA-Seq Kit v3 - Pico Input Mammalian (Takara, cat# 634485) and run on an Illumina NextSeq 2000 (100-bp single-end reads).

### Computational Analyses for RNA Sequencing

For F1 allele-specific samples, reads were aligned to an mm10 N-masked genome generated using SNPsplit^30^, in which SNPs between the C57BL/6 and *Mus musculus castaneus* alleles are replaced with an N nucleotide in the mm10 genome. Alignment was performed using STAR^40^ (v2.7.1a) with options [--alignEndsType EndToEnd --outSAMattributes NH HI NM MD --outSAMtype BAM Unsorted]. Aligned reads were then assigned to a parental genome using SNPsplit^30^ and gene counts were generated on both the diploid N-masked aligned reads and haploid allele-specific aligned reads using featureCounts with options [-p -Q 30 -s 2]^41^. Haploid and diploid count files were input into our EscapeGeneCalculator (https://github.com/Montserrat-Anguera/escapegenecalculator) which identifies escape genes using previously described thresholds^14,15,42^: (1) diploid RPKM > 1, (2) haploid Xi-SRPM > 2, (3) a 95% confidence interval greater than 0 in at least 3 out of 4 replicates. For both F1 allele-specific and TcKO samples, edgeR (v3.32.1) was used to filter and normalize data then DESeq2 (v1.30.1) was used to identify differentially expressed genes with an adjusted p-value ≤ 0.05 and an absolute log2 fold-change ≥ log2(1.5).

### Single-molecule Fluorescent In Situ Hybridization (smFISH) for *Cxcr3* & Image processing

Isolated cells for lung and spleen were suspended in PBS and cytospun onto positively charged slides and permeabilized, fixed with PFA, and stored at -80°C. In preparation for smFISH, slides were rehydrated, then primary *Cxcr3* probes (Integrated DNA Technologies, IDT) were added, heated for 3 minutes at 60°C then incubated overnight at 37°C in a humidified chamber. Slides were washed in 2X SSCT for 5 minutes, 2X SSCT at 60°C for 15 minutes, and twice in room temperature 2X SSCT for 10 minutes. For secondary hybridization, slides were incubated with secondary probes in hybridization buffer and incubated for 2 hours at 37°C in a humidified chamber. Slides were washed as before. Finally, slides were dried, mounted with Gold Antifade Mountant with DAPI, sealed with clear nail polish, and cured for 1 hour prior to imaging.

Acquired images were maximum projected using Fiji ImageJ then analyzed using a MATLAB signal identification algorithm. Briefly, the MATLAB code was developed to: 1) Segment nuclei based on nuclei DAPI signals; 2) ensure each nucleus is unique based on morphology and signal intensities in all channels; and 3) Extract ATTO 565 and ATTO 647N signals, representing the respective exon and intron of *Cxcr3* transcripts. Exon and intron signals were identified based on fitting a Gaussian distribution to signals exceeding an intensity threshold based on the nuclear background signal, and a punctum size threshold. These criteria were validated by manually referencing each identified ATTO 565 and ATTO 647N signal and kept consistent across all experiments. Normalization was done on each cell to avoid experimental variability in puncta determination. Each potential punctum was manually checked to persist across at least two z-stacks (0.54um each) before being marked as either Nascent or Mature *Cxcr3* signal. Mature (ATTO 565 only) and nascent (ATTO 647N only or ATTO 647N overlapping ATTO 565) signals were identified for individual nuclei using our MATLAB algorithm, then manually validated with Fiji Image. Nuclei with two or more Nascent signals that passed the above criteria were counted as ‘biallelic’. Any punctum that overlapped with noise on the 488nm channel was discarded. Chi-square tests were used to determine statistical significance between the TcKO and WT conditions for each cell type in both the lung and the spleen.

### Statistical Analysis

Statistical analyses were performed using GraphPad Prism 11, R Studio, and the indicated bioinformatics packages. Independent two-group comparisons used two-tailed Mann–Whitney U tests. Ordinary two-way ANOVA or two-way repeated-measures ANOVA followed by Holm–Šídák multiple-comparisons tests was used where indicated. Longitudinal body-weight changes were analyzed using mixed-effects models with restricted maximum likelihood (REML) estimation followed by Šídák’s multiple-comparisons tests. Survival was analyzed using the log-rank (Mantel–Cox) test. CUT&RUN, RNA-seq, and *Cxcr3* smFISH statistics are described in their respective Methods sections and figure legends. Data are shown as mean ± SD or SEM, as indicated. Significance was defined as *P* < 0.05 unless otherwise specified. Sample sizes, pooled samples, and biological replication are specified in figure legends.

## Data Availabilty

All sequencing data generated in this study has been deposited to the NCBI GEO database.

## Acknowledgements

We would like to thank the PennVet CHMI for their help with sequencing, C. Hunter, M. May for helpful discussions, and all members from Anguera lab for thoughtful insights during lab meetings. We also thank K. Sarma for pA-MNase used for CUT&RUN experiments, and L. Eisenlohr and M. Hogan for the gift of influenza PR8 viral stocks. This research was supported by NIH R01 AI134834 and R01 AI168047, pilot grants from Penn Vet IIZD & Penn RNA Institute for Innovation, and Lupus Research Alliance Target in Lupus grant (to M.C.A.).

