## Supplemental Figures for "Xist deletion in T cells protects against influenza-induced lung injury"

#### Supplemental Figure Legends

**Supplemental Figure 1. Flu-infected draining lymph node (dLN) T cells exhibit similarly low levels of Xist RNA and H2AK119Ub foci at 5 and 10 dpi.**

CD4$^+$ and CD8$^+$ Naive and AgExp T cells from dLNs of influenza-infected female C57BL/6 mice at 5 and 10 dpi were analyzed using sequential Xist RNA FISH and H2AK119Ub IF. (**A-D**) Representative field images for Xist RNA FISH (top) and IF for H2AK119Ub (bottom) for CD4$^+$ Naive (A), CD4$^+$ AgExp (B), CD8$^+$ Naive (C), and CD8$^+$ AgExp (D) T cells. (**E**) Quantification of the percentage of nuclei with Xist RNA signals, consisting of clouds, dispersed clusters, or pinpoints. (**F**) Quantification of H2AK119Ub foci for each T-cell subset. For each biological replicate and T-cell subset, 21–210 nuclei were quantified for Xist RNA signals and 13–216 nuclei for H2AK119Ub foci via IF. For 5 dpi, each data point represents T cells from $n = 2$ mice pooled together; for 10 dpi, each data point represents one mouse. Arrows indicate Xist RNA signals that colocalize with a H2AK119Ub focus. Scale bars, 10 μm. Data are shown as mean ± SEM. CD4$^+$ and CD8$^+$ T-cell subsets were analyzed separately by ordinary two-way ANOVA with time point and T-cell subset as factors, followed by Holm–Šídák multiple-comparisons tests comparing 5 vs. 10 dpi within each T-cell subset.

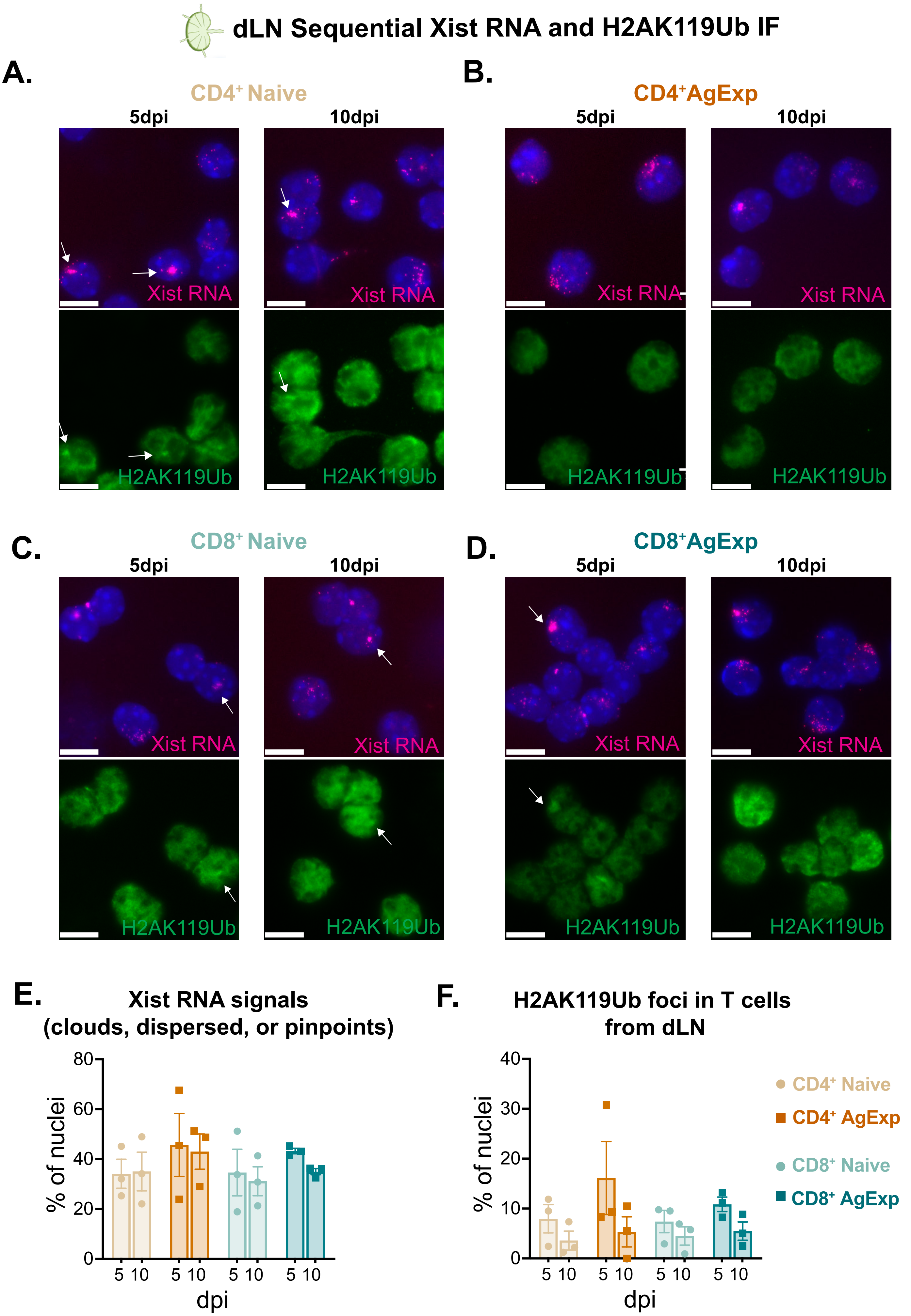


Supplemental Figure 1

**Supplemental Figure 2. Flu-infected splenic T cells exhibit similarly low levels of Xist RNA and H2AK119Ub foci at 5 and 10 dpi.**

$CD4^+$ and $CD8^+$ Naive and AgExp T cells from spleens of influenza-infected female C57BL/6 mice at 5 and 10 dpi were analyzed using sequential Xist RNA FISH and H2AK119Ub IF. (**A-D**) Representative field images for Xist RNA FISH (top) and IF for H2AK119Ub (bottom) for $CD4^+$ Naive (A), $CD4^+$ AgExp (B), $CD8^+$ Naive (C), and $CD8^+$ AgExp (D) T cells. (**E**) Quantification of the percentage of nuclei with Xist RNA signals, consisting of clouds, dispersed clusters, or pinpoints. (**F**) Quantification of H2AK119Ub foci for each T-cell subset. Arrows indicate Xist RNA signals that colocalize with a H2AK119Ub focus. Scale bars, 10 µm. For each biological replicate and T-cell subset, 61–148 nuclei were evaluated for Xist RNA and 39–143 nuclei for H2AK119Ub IF. For 5 dpi, each data point represents T cells from $n$ = 2 mice pooled together; for 10 dpi, each data point represents one mouse. Data are shown as mean ± SEM. $CD4^+$ and $CD8^+$ T-cell subsets were analyzed separately by ordinary two-way ANOVA with time point and T-cell subset as factors, followed by Holm–Šídák multiple-comparisons tests comparing 5 vs. 10 dpi within each T-cell subset. ns, not significant.

**Spleen Sequential Xist RNA FISH and H2AK119Ub IF**

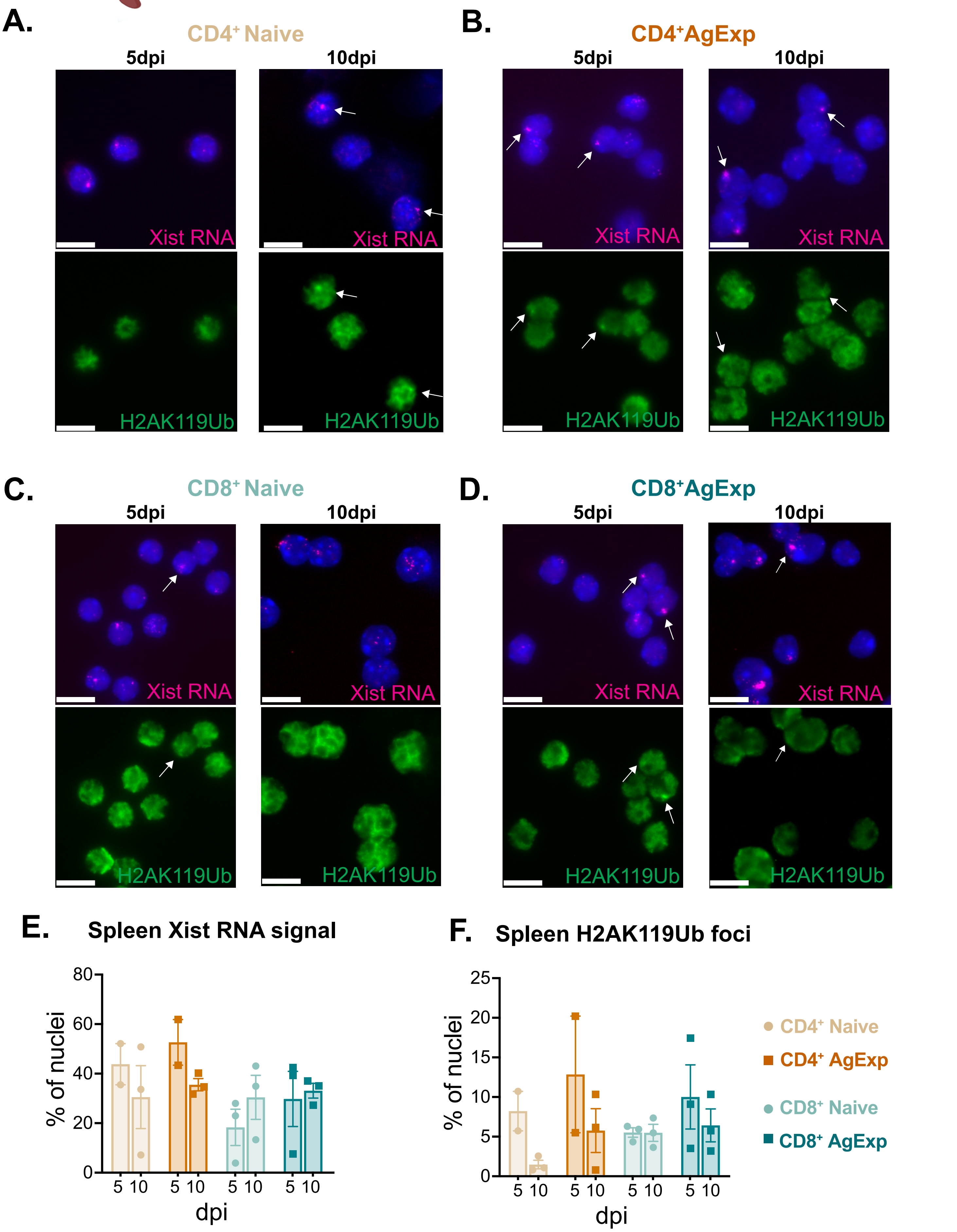


**Supplemental Figure 2**

**Supplemental Figure 3. Allelic profiling of heterochromatic marks H2AK119Ub and H3K27me3 across Chromosome 13 in flu-infected T cell subsets.**

(**A**) Genome browser tracks of CPM normalized reads for H2AK119Ub and H3K27me3 at the *Hox* gene cluster and the neighboring expressed gene *Skap2* in CD4$^+$ Naive and AgExp and CD8$^+$ lung T cells at 5 and 10 dpi. Colors (purple and blue respectively) represent the paternal *castaneus* allele and black represents the maternal C57BL/6 allele. Gray bars indicate unmappable regions. (**B**) Genome browser tracks of CPM normalized reads for H2AK119Ub and H3K27me3 for Chromosome 13. (**C**) Mean d-score [(Xireads/(Xireads+Xareads)) – 0.5] at genic and intergenic regions on chromosome 13 between 5 and 10 dpi in CD4$^+$ Naive T cells. Genic regions are defined as +1 kb from the transcription start site (TSS) to the transcription end site (TES) and intergenic regions are defined as 10 kb bins not overlapping genes or their promoters. Adjusted *P*-values were calculated using a Kruskal-Wallis test followed by a Dunn's test with Benjamini-Hochberg correction for pairwise comparisons. (**D-E**) Mean d-score at genic and intergenic regions on chromosome 13 between 5 and 10 dpi for H2AK119Ub (D) and H3K27me3 (E) in CD4$^+$ AgExp lung T cells. (**F-G**) Mean d-score at genic and intergenic regions on chromosome 13 between 5 and 10 dpi for H2AK119Ub (F) and H3K27me3 (G) in CD8$^+$ Naive lung T cells. (**H-I**) Mean d-score at genic and intergenic regions on chromosome 13 between 5 and 10 dpi for H2AK119Ub (H) and H3K27me3 (I) in CD8$^+$ AgExp lung T cells. (**J-M**) Normalized read counts for H2AK119Ub (purple) and H3K27me3 (blue) in 10kb bins across the Cast chromosome 13 in 5 dpi (top) and 10 dpi (bottom) CD4$^+$ Naive lung T cells (J), CD4$^+$ AgExp lung T cells (K), CD8$^+$ Naive lung T cells (L), and CD8$^+$ AgExp lung T cells (M). Gray bars indicate unmappable regions, and pink bar represents the *Xist* locus.

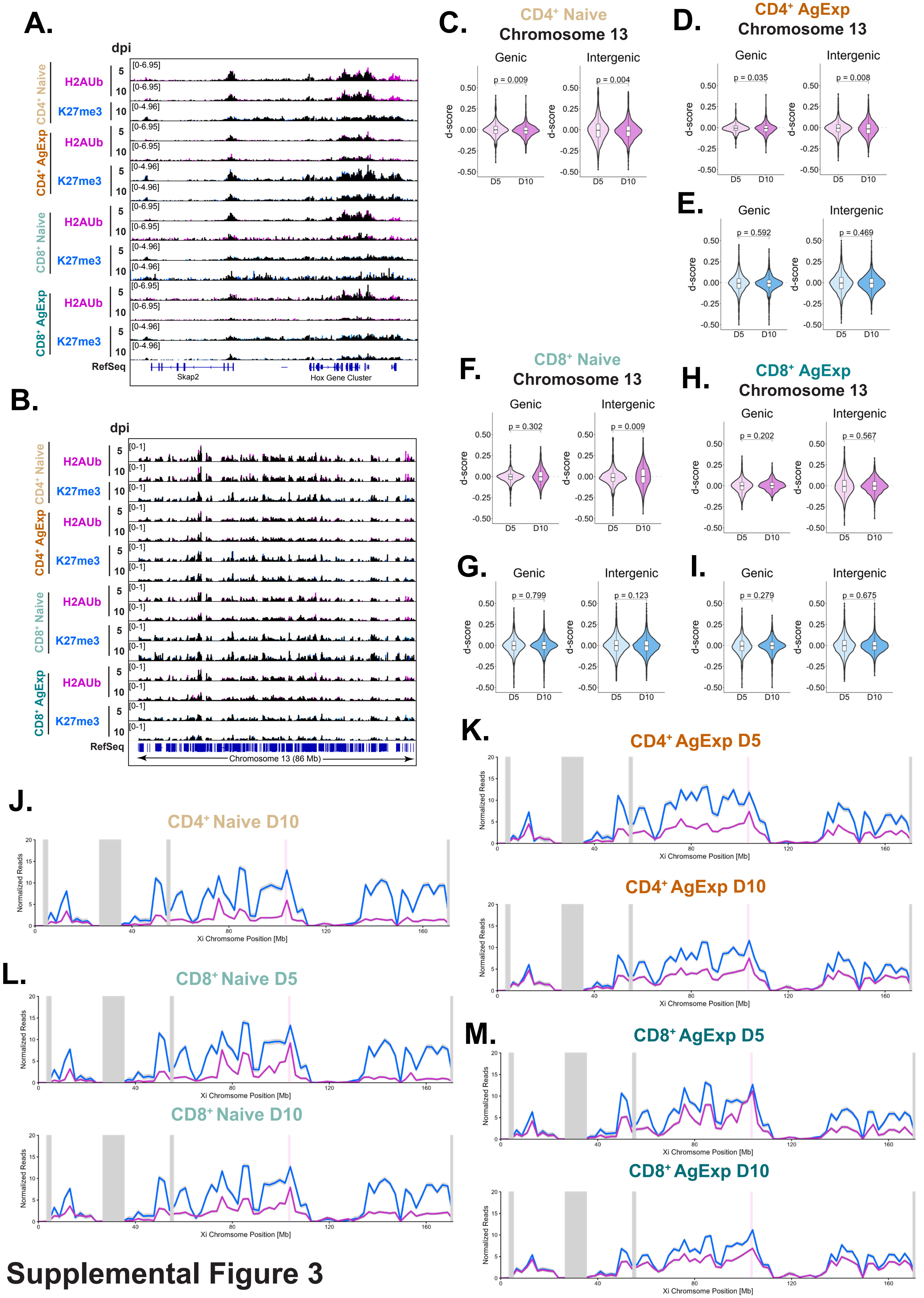


**Supplemental Figure 3**

**Supplemental Figure 4. Differential gene expression analyses of infected lung T cell subsets.**

(**A**) Principal component analysis (PCA) of RNA-seq profiles from lung $CD4^+$ and $CD8^+$ Naive and AgExp T cells isolated from female F1 mus x cast mice at 10 dpi. (**B**) Heatmap of the top 50 genome-wide differentially expressed genes (DEGs), ranked by adjusted *P* value, from each $CD4^+$ and $CD8^+$ AgExp vs. Naive T cell comparison at 10 dpi. (**C**) Allele-specific expression [SNP-reads per million (SRPM)] from the maternal autosomes (black) and paternal autosomes (colors) in $CD4^+$ and $CD8^+$ Naive and AgExp T cells isolated from the lung 10 dpi. (**D**) *Xist* expression in $CD4^+$ and $CD8^+$ Naive and AgExp T cells from infected lungs, shown as SRPM. Each data point represents one mouse, and data are shown as mean ± SD. *Xist* expression was analyzed by two-way repeated-measures ANOVA followed by Holm–Šídák multiple-comparisons tests. (**E**) Normalized Xi enrichment of H2AK119Ub and H3K27me3 at all identified XCI escape genes in $CD4^+$ and $CD8^+$ Naive or AgExp T cells from at 10 dpi [+1 kb from transcription start site (TSS) to the transcription end site (TES)]. Each column represents 1 biological replicate. The T cell subset(s) in which XCI escape was identified (**Supplemental File 1**) is denoted by colored boxes on the left.

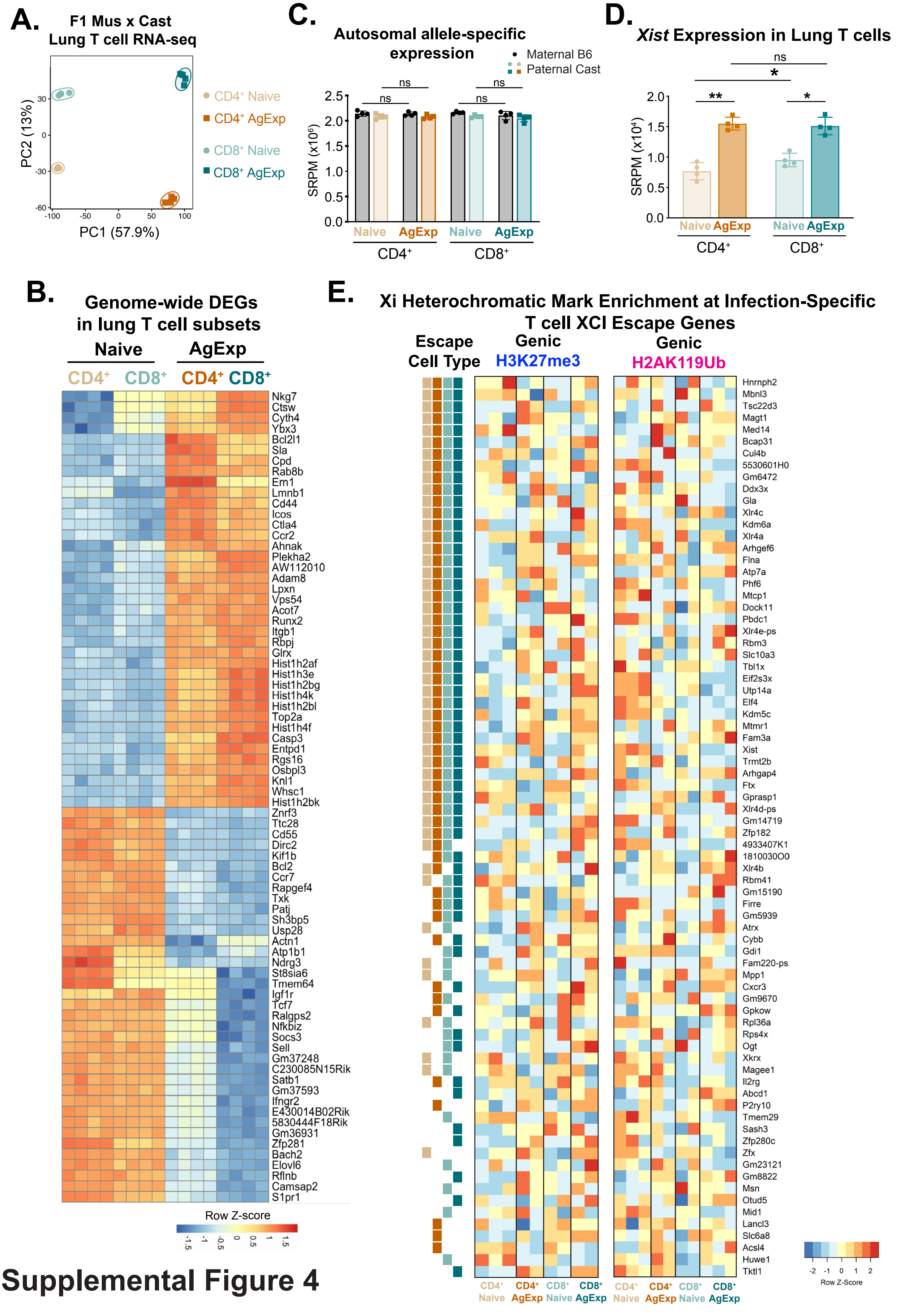


Supplemental Figure 4

**Supplemental Figure 5. Validation of Xist RNA deletion in female Xist TcKO CD3$^+$ T cells and quantification of splenic T cell compartment in flu-infected WT and Xist TcKO mice.**

(**A–B**) Splenic CD3$^+$ T cells from naive female Xist WT and Xist TcKO mice were stimulated with anti-CD3/CD28 for 72 hrs then used for sequential Xist RNA FISH and IF. Representative field images and quantification of Xist RNA signals from Xist RNA FISH (A) and focal H3K27me3 localization by immunofluorescence (IF) (B) are shown. Each data point represents one mouse. Scale bars, 10 µm. (**C**) Survival rates for female WT (n=10) and Xist TcKO (n=7) mice over time. Survival curves were compared using the log-rank (Mantel–Cox) test; ns, not significant. (**D**) Frequency and number of lung CD19$^+$ B cells at 10 dpi. (**E**) Influenza-specific hemagglutination inhibition (HAI) antibody titers in serum at 30 dpi. Data were pooled from three independent experiments with 3–8 mice per group. (**F**) Spleen weight and cellularity at 10 dpi. Data were pooled from three independent experiments with 3–5 mice per group. (**G–M**) Flow-cytometric analysis of splenic T cell populations at 10 dpi, showing frequency and cell/mg of spleen for Foxp3$^+$ Treg cells (G), total CD4$^+$ T cells (H), CD4$^+$ Naive T cells (I), CD4$^+$ AgExp T cells (J), total CD8$^+$ T cells (K), CD8$^+$ Naive T cells (L), and CD8$^+$ AgExp T cells (M). Each data point represents one mouse, and Xist WT and Xist TcKO groups were compared using two-tailed Mann–Whitney tests. Data are shown as mean ± SEM. *$P < 0.05$, **$P < 0.01$; ns, not significant.

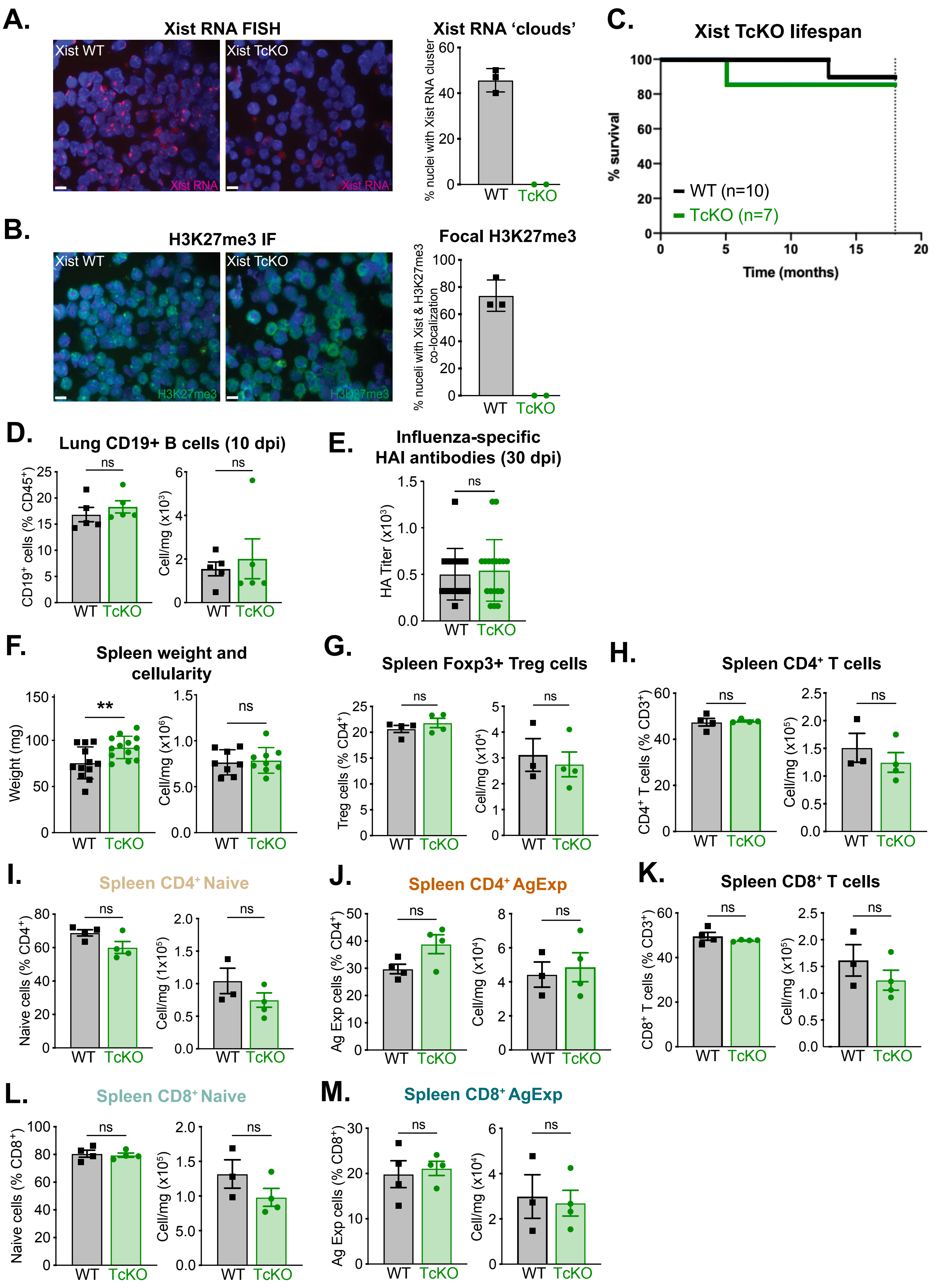


**Supplemental Figure S5**

**Supplemental Figure 6. SmFISH analyses for quantification of monoallelic and biallelic *Cxcr3* expression in splenic Cxcr3$^+$ AgExp T cells following influenza infection of WT and Xist TcKO mice.**

(**A–B**) Representative smFISH images for allelic expression of nascent *Cxcr3* expression showing signals from intronic Cxcr3 Cy3-labeled probes for CD4$^+$ (A) and CD8$^+$ (B) AgExp Cxcr3$^+$ T cells from spleens of infected mice at 10 dpi. Scale bars, 10 μm. Staining with exonic *Cxcr3* probes was performed (not shown) and used for MATLAB puncta quantification to call nascent pinpoint transcription. (**C–D**) Percentage of biallelic *Cxcr3*-transcribing cells for CD4$^+$ (C) and CD8$^+$ (D) Cxcr3$^+$ AgExp T cells from infected lungs. For WT samples, each data point represents one mouse (n = 3); for TcKO samples, one data point represents T cells from one mouse and the other data point corresponds to T cells pooled together from n=3 infected mice. (**E-F**) Percentage of biallelic *Cxcr3*-transcribing cells for CD4$^+$ (C) and CD8$^+$ (D) Cxcr3$^+$ AgExp T cells from spleen. For spleen analyses, each data point represents one mouse (WT, n = 3; TcKO, n = 4). Data are representative of two independent experiments with 3–4 mice per group. WT and TcKO groups were compared using two-tailed Mann–Whitney tests. Data are shown as mean ± SD. *$P$ < 0.05; ns, not significant.

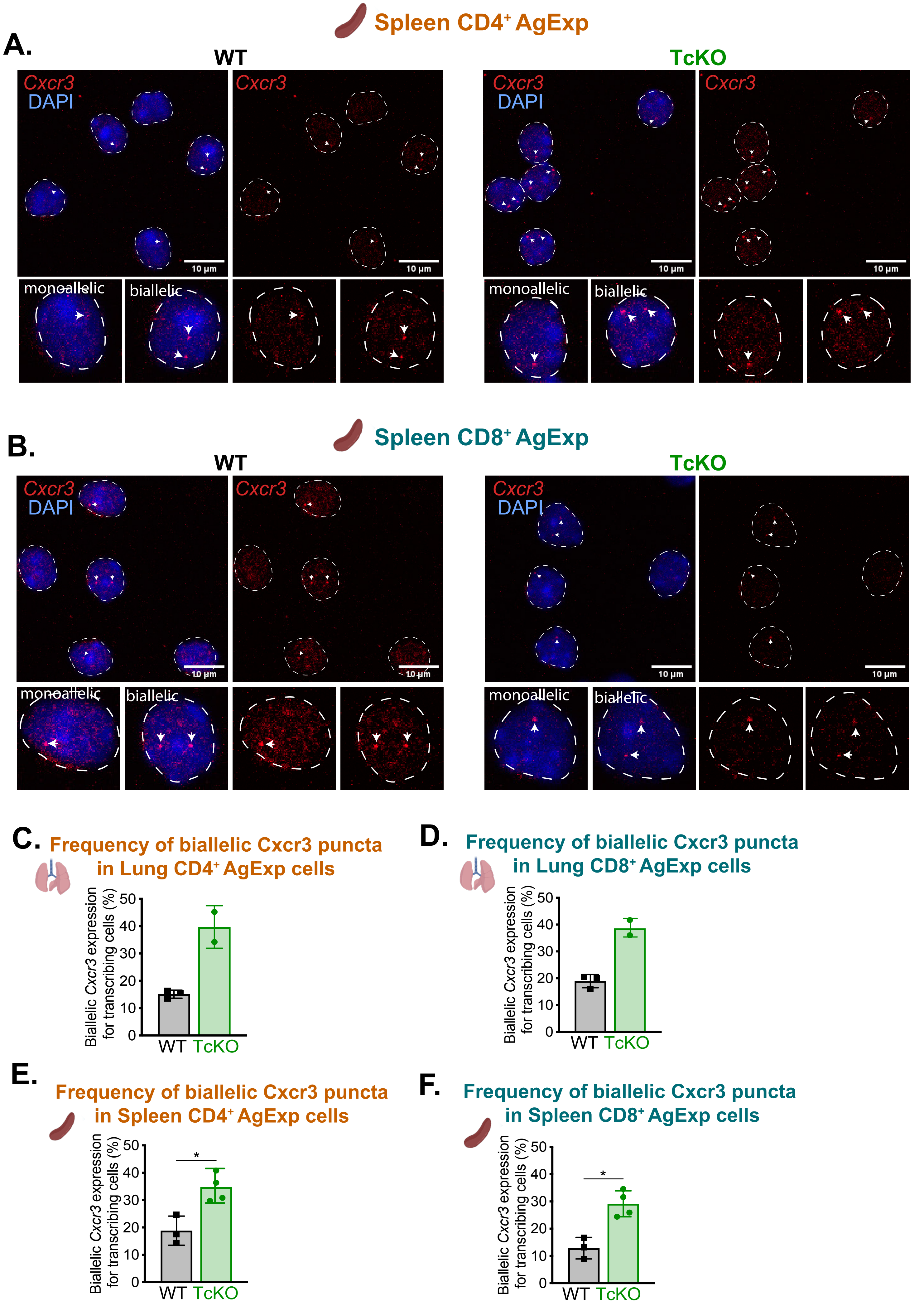


**Supplemental Figure S6**

**Supplemental Figure 7. T cell-specific *Xist* deletion enhances Cxcr3 protein levels in the splenic T cell compartment following influenza infection.**

**(A, D)** Representative UMAP plots of splenic $CD4^+$ (A) and $CD8^+$ (D) T cells from Xist WT and Xist TcKO female mice at 10 dpi, showing Naive, AgExp, and Treg populations and expression of the indicated markers, including Cxcr3. **(B–C, E–F)** Flow cytometric quantification of surface Cxcr3 protein within splenic $CD4^+$ Naive (B), $CD4^+$ AgExp (C), $CD8^+$ Naive (E), and $CD8^+$ AgExp (F) T cell populations from infected mice. The frequency and number of $Cxcr3^+$ cells/mg of spleen is shown. Data are representative of two independent experiments with 3–4 mice per group. Each data point represents one mouse. Data are shown as mean ± SEM. Xist WT and Xist TcKO groups were compared using two-tailed Mann–Whitney tests. *$P < 0.05$; ns, not significant.

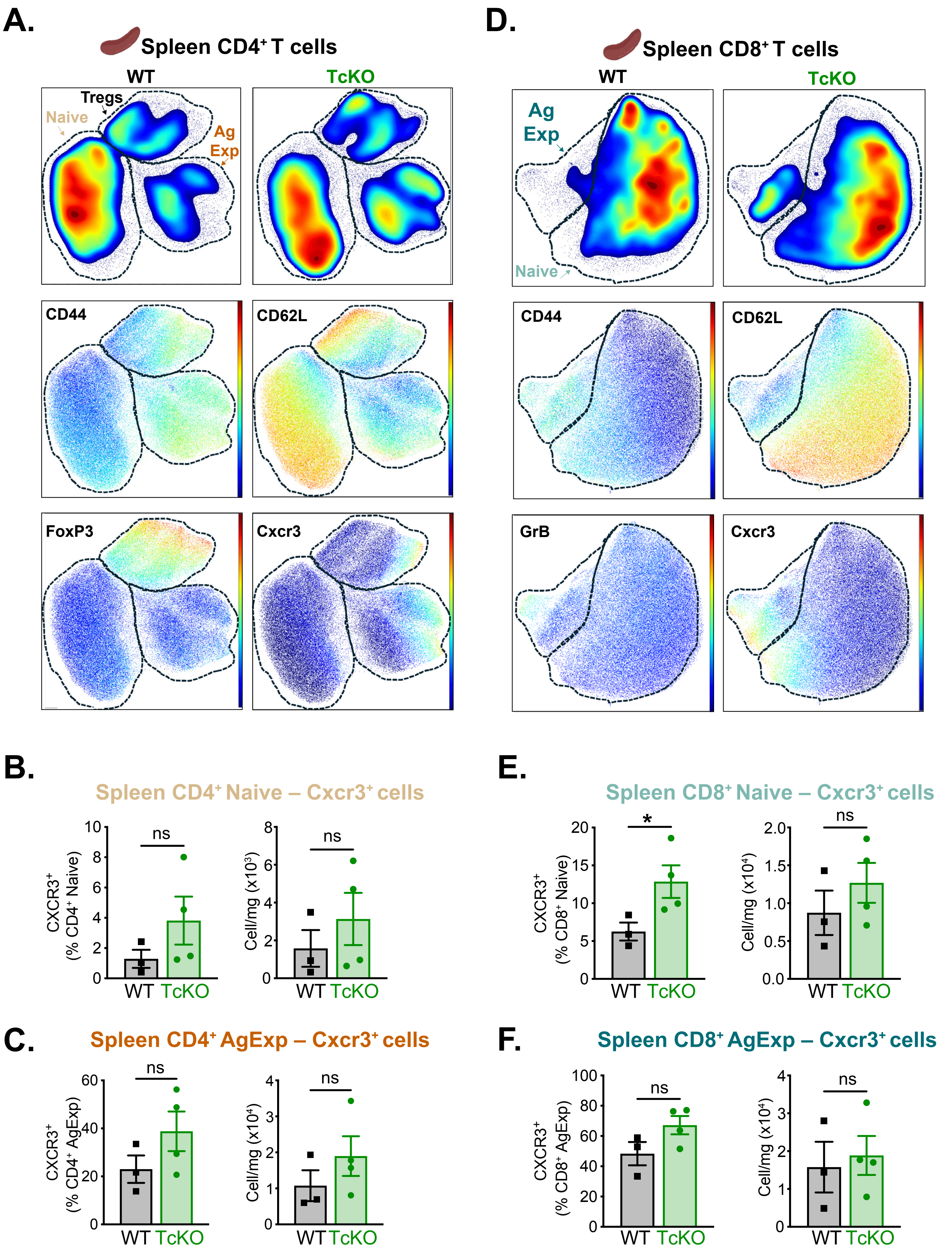


**Supplemental Figure S7**

**Supplemental Figure 8. *Xist* deletion enhances the cytokine-producing potential of AgExp T cells from the spleens of infected mice.**

(**A**) Experimental design for *ex vivo* T-cell stimulation using spleens from infected mice. Female Xist WT and Xist TcKO mice were infected with influenza PR8, and dissociated splenocytes at 10 dpi were stimulated with PMA/Ionomycin for 4h, followed by intracellular cytokine staining. (**B,C**) Representative UMAP plots of splenic $CD4^+$ (B) and $CD8^+$ (C) T cells from WT and Xist TcKO mice following *ex vivo* stimulation, showing Naive, AgExp, and Treg populations (for $CD4^+$ cells). Expression levels of IFN-γ, TNF, CD40L (B), and GrB are shown overlaid on UMAP plots. (**D**) Frequencies of $Cxcr3^+$, $Cxcr3^+IFN\text{-}\gamma^+$, and $IFN\text{-}\gamma^+TNF^+$ T cells for $CD4^+$ Naive and AgExp T cells after *ex vivo* stimulation of splenocytes. (**E**) Frequencies of $Cxcr3^+$, $Cxcr3^+IFN\text{-}\gamma^+$, and $IFN\text{-}\gamma^+TNF^+$ T cells for $CD8^+$ Naive and AgExp T cells after *ex vivo* stimulation of splenocytes. Data are representative of two independent experiments with n = 4 mice per group. Each data point represents one mouse. Data are shown as mean ± SEM. Xist WT and Xist TcKO groups were compared using two-tailed Mann–Whitney tests. *$P < 0.05$; ns, not significant. (**F**) Frequencies of $CD107a^+$ cells within $CD4^+$ and $CD8^+$ Naive and AgExp T cells from lungs of infected mice (10 dpi). (**G**) Frequencies of $Cxcr3^+CD107a^+$ cells within $CD4^+$ and $CD8^+$ Naive and AgExp T cells from lungs of infected mice (10 dpi). Data for (F-G) were pooled from two independent experiments with 3–5 mice/group, and WT and TcKO groups were compared using two-tailed Mann–Whitney test. Data are shown as mean ± SD.

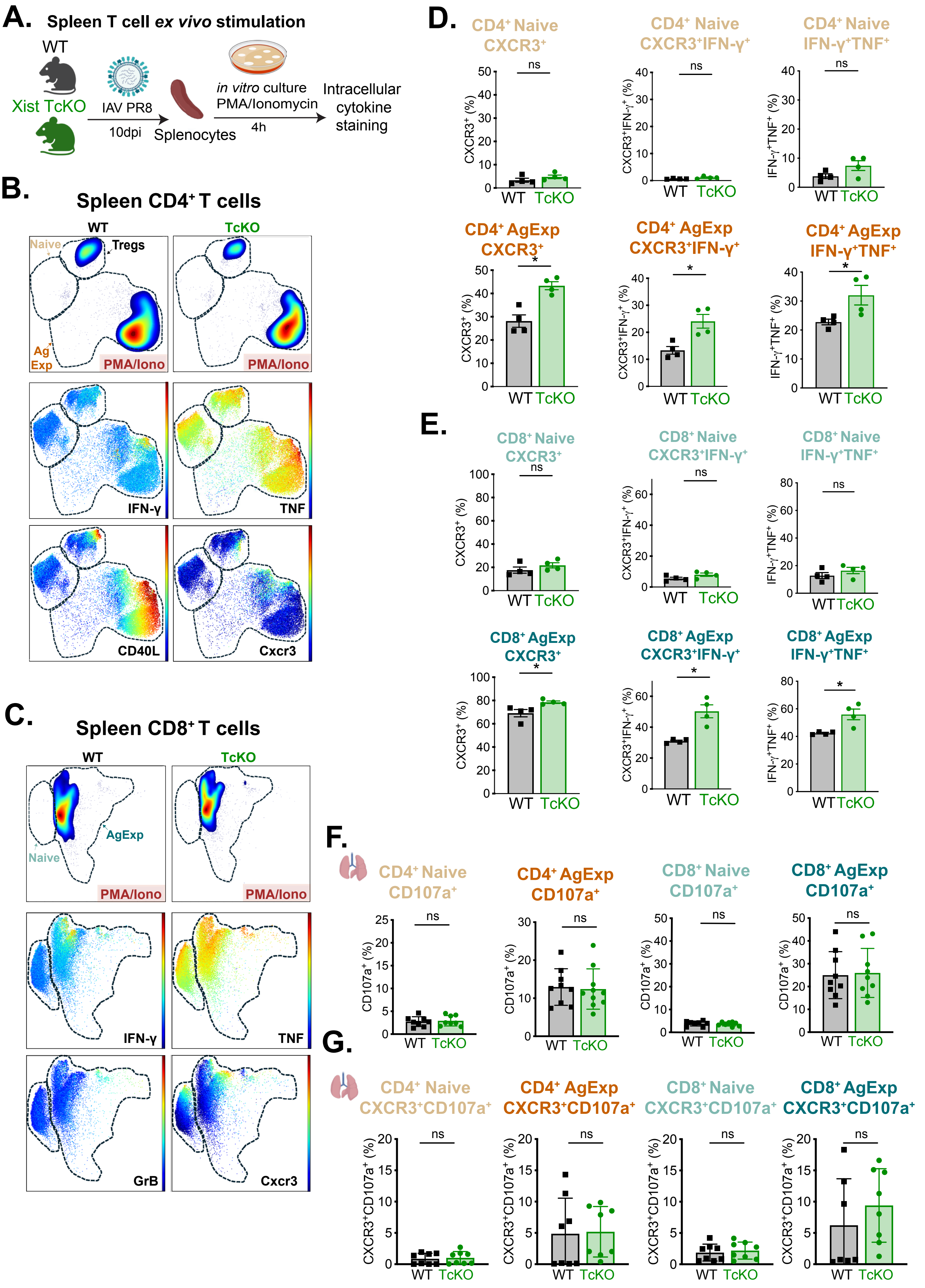


### Supplemental Figure S8
